# Multiple weak brakes act in concert to regulate STIM1 and control store-operated calcium entry

**DOI:** 10.1101/2025.05.20.655205

**Authors:** Ruoyi Qiu, Richard S. Lewis

**Affiliations:** Department of Molecular and Cellular Physiology, Stanford University School of Medicine, Stanford, CA 94305

## Abstract

The ER Ca^2+^ sensor STIM1 evokes store-operated Ca^2+^ entry through the interaction of its cytosolic CRAC activation domain (CAD) with the plasma membrane Ca^2+^ channel Orai1 following ER Ca^2+^ depletion. To avoid pathophysiological effects, STIM1 must respond precisely and reversibly to changes in ER Ca^2+^ content while maintaining a low basal activity under resting, ER-replete conditions. Here we combine single-molecule FRET measurements of full-length dimeric STIM1 in lipid membranes with an AlphaFold2 structural model to describe the structure and regulation of the resting state. We show that STIM1 activity is controlled by the combined operation of four relatively weak restraints, or brakes. The Ca^2+^-bound EF-SAM luminal domain acts as a steric restraint to inhibit spontaneous activity. In the cytosolic region, the domain-swapped interaction and alignment of CC1α1 with CC3 of CAD positions the apex of CAD next to the ER membrane, where electrostatic lipid-protein interactions further stabilize the inactive conformation. A fourth brake is created by hydrophobic and electrostatic interactions of the two CC1α2/3 domains attached to the base of CAD. Disruption of any one of these brakes leads to STIM1 activation, showing that the concerted action of these relatively weak restraints is required to prevent spontaneous activity in resting cells with full ER Ca^2+^ stores, while enabling reversible activation in response to changes in store content.

## INTRODUCTION

Store-operated Ca^2+^ entry (SOCE) is a nearly ubiquitous Ca^2+^ entry pathway activated by cell-surface receptors that release ER Ca^2+^, typically through the activation of IP_3_ receptors in the endoplasmic reticulum (ER) (Prakriya and Lewis, 2015). SOCE is essential for the immune response, as well as ensuring proper muscle development and function, exocrine secretion, and many other functions, and naturally-occurring mutations have been linked to a spectrum of human health disorders (Vaeth et al., 2020; Korshunov and Prakriya, 2025; Protasi et al., 2023; Concepcion and Feske, 2017). GOF mutations lead to tubular aggregate myopathy, Stormorken syndrome, and York platelet syndrome, while LOF mutations cause immunodeficiency, muscular hypotonia, autoimmunity, among other disorders, demonstrating the critical importance of regulation and reversibility of SOCE (Feske, 2019).

The ER protein STIM1 is the principal regulator of SOCE and operates by controlling access to two critical cytosolic domains: the polybasic domain and CAD (CRAC activation domain), also known as SOAR (STIM1-Orai activating region) (Park et al., 2009; Yuan et al., 2009) (**Fig. 1A**). Depletion of ER Ca^2+^ is sensed by the luminal cEF hand of STIM1, triggering a large conformational change that (1) exposes the polybasic domain so it can bind phosphoinositide plasma membrane (PM) lipids and promote accumulation at ER-PM junctions, and (2) extends the CAD domain towards the PM where it binds and activates the CRAC channel Orai1. STIM1 acts as a switch: once activated, all subsequent steps leading to SOCE occur through a passive diffusion trap mechanism, and the sequence is reversed upon refilling of ER Ca^2+^ stores (Prakriya and Lewis, 2015).

**Figure 1.**
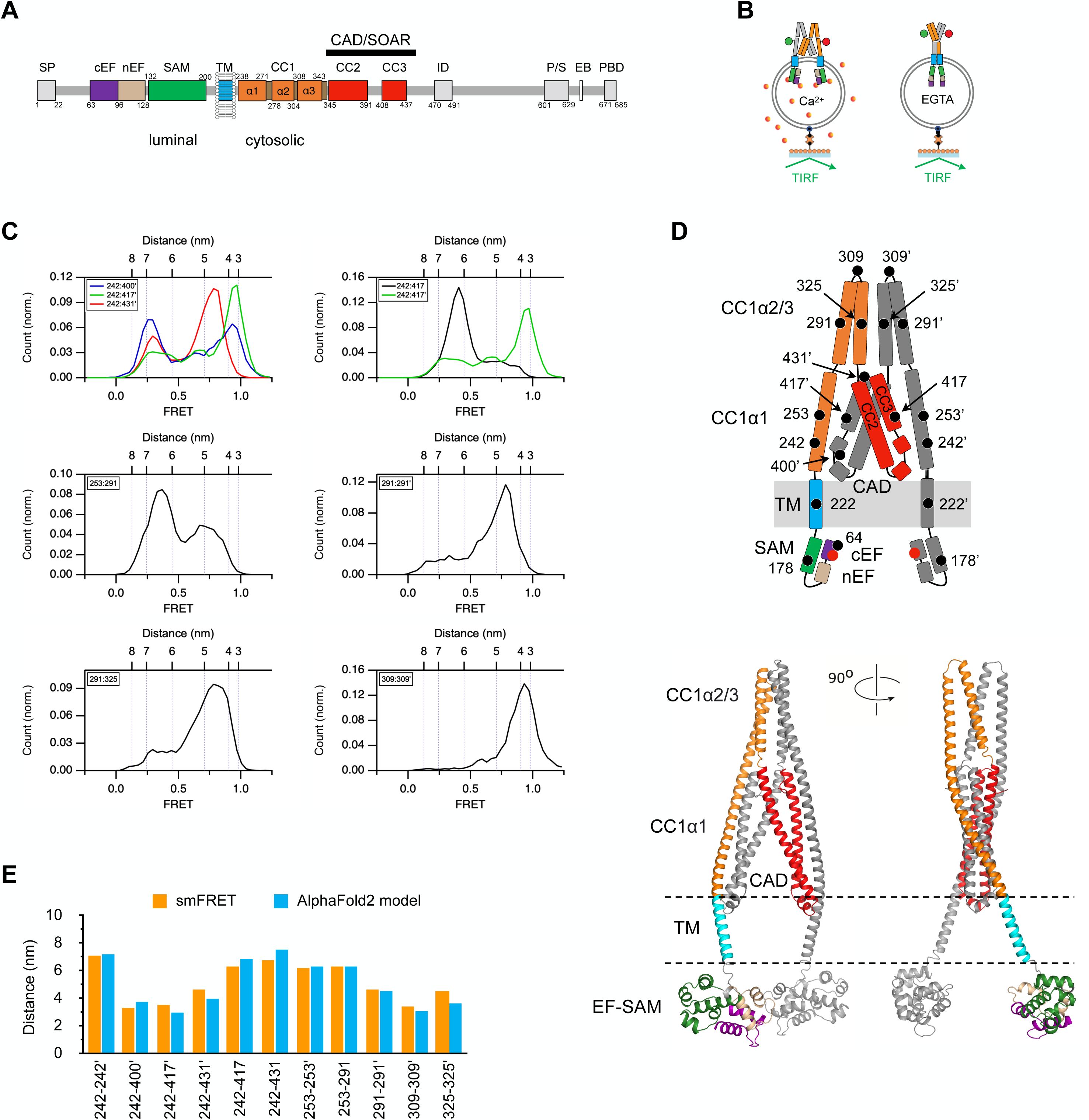
Conformation of STIM1 in the Ca^2+^-bound inactive state. A. Diagram of STIM1 functional domains: SP, signal peptide; cEF, canonical EF hand; nEF, non-canonical EF hand; SAM, sterile alpha motif; TM, transmembrane domain; CC1-3, coiled-coil 1-3; ID, inactivation domain; P/S, proline/serine-rich region; EB, EB1 binding domain; PBD, polybasic domain; CAD, CRAC activation domain; SOAR, STIM-Orai activation region. Residue numbers for each domain are indicated. B. flSTIM1 labeled with donor and acceptor dyes is incorporated into liposomes in the presence of 2 mM Ca^2+^ or 0.5 mM EGTA, and attached to coverglass substrate via a biotin-streptavidin linker. The donor dye is excited in TIRF mode, and donor and acceptor dye emissions are used to calculate FRET. C. Selected smFRET amplitude histograms in 2 mM Ca^2+^ supporting key features of the structure in D. 242:400’ (n=188 molecules), 242:417’ (n=190), 242:431’ (n=189), 242:417 (n=291), 253:291 (n=254), 291:291’ (n=185), 291:325 (n=202), 309:309’ (n=76). Distances are calculated from FRET values using an R_0_ value of 5.8 nm. D. Diagram of STIM1 topology (top) and an AlphaFold2 model (bottom) consistent with smFRET measurements. For clarity, both show only the structure from the cEF hand to the N-terminal end of CAD (residues 63 to 444). Colors correspond to the domain diagram in A, and bound Ca^2+^ ions are indicated by the red dots. E. A comparison of inter-dye distances within the cytosolic domain derived from smFRET measurements and the AlphaFold2 model.

To avoid GOF pathologies such as Stormorken syndrome and tubular aggregate myopathy, STIM1 must be held inactive at normal resting levels of ER Ca^2+^ (∼0.5 mM). Multiple lines of evidence support a critical role of the CC1α1 domain and the CC3 domain of CAD in enforcing the inactive state; this is often referred to as the ‘CC1 clamp.’ The isolated CC1α1 and CC3 domains interact directly as detected by FRET (Fahrner et al., 2014), and in a split-STIM assay in which an ER-bound STIM fragment containing the CC1α1 domain binds to a soluble cytosolic

CAD fragment and releases it upon ER Ca^2+^ depletion (Ma et al., 2015, 2020; Shrestha et al., 2022; Korzeniowski et al., 2017; Ma et al., 2017). Multiple studies have identified mutations in CC1α1 and CC3 that produce constitutive ([Ca^2+^]_ER_-independent) SOCE and affect CC1α1-CC3 physical interactions (Muik et al., 2009, 2011; Shrestha et al., 2022), but the actual binding interface has not been identified. CC1α3 has also been proposed as a brake based on a quadruple mutation (4EA; E318A/319A/320A/322A) that activates STIM1, although the mechanism remains unclear (Korzeniowski et al., 2010; Yu et al., 2013). These studies raise questions about the number and structural bases of brakes that keep STIM1 in cells inactive under resting conditions.

Structural analysis with x-ray crystallography or cryo-EM could provide deeper insights into the mechanisms of STIM1 regulation, but the protein’s high flexibility has thus far thwarted attempts to determine the structure of full-length STIM1 (flSTIM1). What structural information we have derives from studies of STIM1 fragments: a crystal structure of CAD (Yang et al., 2012) and NMR structures of the Ca^2+^-bound EF-SAM domain (Stathopulos et al., 2008), the CC1 monomer (Rathner et al., 2020), and a CC1α3-CC2 fragment (Stathopulos et al., 2013). While these partial structures can provide much useful information, it is difficult to know a priori whether and under what conditions they exist in the full-length protein and if they do, what their functions are. Tests based on the effects of mutagenesis generally support their proposed functions as brakes, with the caveat that several sites are known to participate in more than one conformational state. For example, L251 negatively regulates STIM1 through the CC1α1-CC3 clamp in the resting state but positively regulates STIM1 after store depletion by helping form an extended CC1 coiled-coil (Zhou et al., 2013; Hirve et al., 2018). The 4EA mutation inhibits the resting STIM1 state by destabilizing the CC1α2/3 brake (this paper), but may also promote Orai1 activity by stabilizing a CC1α3-CC2 structure proposed to bind to Orai1 (Stathopulos et al., 2013). The repurposing of these regulatory domains in different states makes it difficult to infer the role of any particular STIM1 conformation or interaction from the effects of mutations on Orai1 activity.

To gain insight into the organization of the cytosolic domain in the resting state, we initially studied a cytosolic domain fragment of STIM1 (ctSTIM1; aa 233-685) with single-molecule Förster resonance energy transfer (smFRET) (van Dorp et al., 2021). smFRET gives information about intramolecular distances as well as conformational dynamics and is thus particularly well suited to study flexible proteins like STIM1. We found that in ctSTIM1, CC1α1 associates with CAD in a domain-swapped configuration with an orientation predicted to position the CAD apex, the region thought to bind to Orai1 (Butorac et al., 2019), next to the ER membrane, and with CC1α2-CC1α3 hairpin structures pointing away from the base of CAD (van Dorp et al., 2021).

However, while ctSTIM1 is a model for the inactive conformation of the cytosolic domain, the absence of the transmembrane and luminal EF hands and SAM domains limits its usefulness for mechanistic studies of STIM1 activation. Specifically, the lack of membrane attachment is likely to alter the forces acting on the cytosolic domain to control CC1-CAD interactions, while the absence of luminal domains precludes studies of activation by Ca^2+^ removal.

Here we have developed and applied a new system to study the regulation of full-length STIM1 (flSTIM1) embedded in artificial membranes. Using smFRET-derived intramolecular distance mapping, we identified an AlphaFold2 model that correctly predicts side-chain interactions underlying the CC1α1-CC3 brake, pinpointing the orientation of CAD relative to the ER membrane in the inactive state and revealing a functional interaction of CAD with the ER membrane. The model structure also defines a novel brake arising from intersubunit interactions between CC1α2/3 helices. Together, these studies offer a new understanding of the multiple intramolecular forces that act together to prevent pathological activation of STIM1 and SOCE in resting cells while preserving the ability to become reversibly activated by changes in ER [Ca^2+^].

## RESULTS

### flSTIM1 resting state structure under Ca^2+^-saturated conditions

To prepare flSTIM1 for dye labeling, we first replaced all of the endogenous cysteines (C49, C56, C227, and C437) with serines. These substitutions did not significantly affect STIM1 activity as indicated by a normal resting [Ca^2+^]_i_ and the amplitude of thapsigargin-induced SOCE (**Supplementary** Fig. 1; Methods). After introducing new cysteines at selected sites, purified flSTIM1 was labeled with donor and acceptor fluorophores, reconstitituted in liposomes in the presence of 2 mM Ca^2+^ or 0.5 mM EGTA, and attached to coverslips for smFRET measurements by TIRF microscopy (**Fig. 1B**). Unless otherwise noted, all data in this study were collected in the presence of 2 mM Ca^2+^, which is expected to saturate the luminal binding sites and generate the inactive structure, given the EF-hand affinity of ∼200 µM (Luik et al., 2008; Gudlur et al., 2018; Stathopulos et al., 2006; Brandman et al., 2007).

smFRET values were collected from 15 dye pairs positioned throughout CC1 and CAD, as summarized in normalized amplitude histograms in **Fig. 1C** and **Supplementary** Fig. 2. Peak FRET values were converted to distances (**Supplementary Table 1**) to constrain a coarse-grained model of the STIM1 resting conformation (**Fig. 1D**), revealing several major features (we use ‘:’ throughout to denote a pair of sites in the dimer and ‘’’ to indicate a site on the adjacent subunit). Distances from 242:400’, 242:417’, and 242:431’ dye pairs indicate that CC1α1 is parallel to CC3 in CAD, and the result that 242 is closer to 417’ than 417 indicates a domain-swapped configuration. This association of CC1α1 with CC3 orients the apex of CAD to face the membrane and keeps the TM and luminal SAM domains well separated. Multiple smFRET-derived distances suggest that the CC1α2 and CC1α3 domains form hairpin structures (291:325) that are closely associated (291:291’, 309:309’, 325:325’) and point away from the base of CAD (253:291). This arrangement of the cytosolic region and corresponding smFRET values are similar to those of the soluble cytosolic fragment of STIM1 (ctSTIM1) we described in a previous smFRET study (van Dorp et al., 2021) (**Supplementary Table 2**), although there are important differences in dynamics stemming from the presence of membrane anchors in the flSTIM1 structure (described below).

To gain more insight into the resting structure in saturating Ca^2+^ and the intramolecular contacts that stabilize it, we generated a series of AlphaFold2 models for the flSTIM1 dimer. Out of the top five models (**Supplementary** Fig. 3), one mirrored the coarse-grained model and was highly consistent with the smFRET data in the cytosolic region (**Fig. 1D**). Within this region, the inter-dye distances simulated on the AlphaFold2 model agree closely (within ± 0.9 nm) with distances calculated from smFRET (**Fig. 1E**). While the model needs further refinement for the TM and luminal domains (see Methods), it proved to be quite useful in revealing important interaction sites within the cytosolic domain, as described below.

The stability of the resting conformation shown in **Fig. 1D** is shown by the large area under the predominant FRET peaks at multiple dye locations (**Fig. 1C**). For convenience, we selected the smFRET measurements of the 242:242’ pair to quantify occupancy of the resting state, because the high probability (70%) of the lowest FRET state in saturating Ca^2+^ (0.28) fell to 25% in EGTA as the distribution shifted to higher FRET states (**Fig. 2A, B**). Below, we quantify the 242:242’ low-FRET probability to assess the function of intramolecular restraints (‘brakes’) that stabilize the resting state under conditions of saturating Ca^2+^.

**Figure 2.**
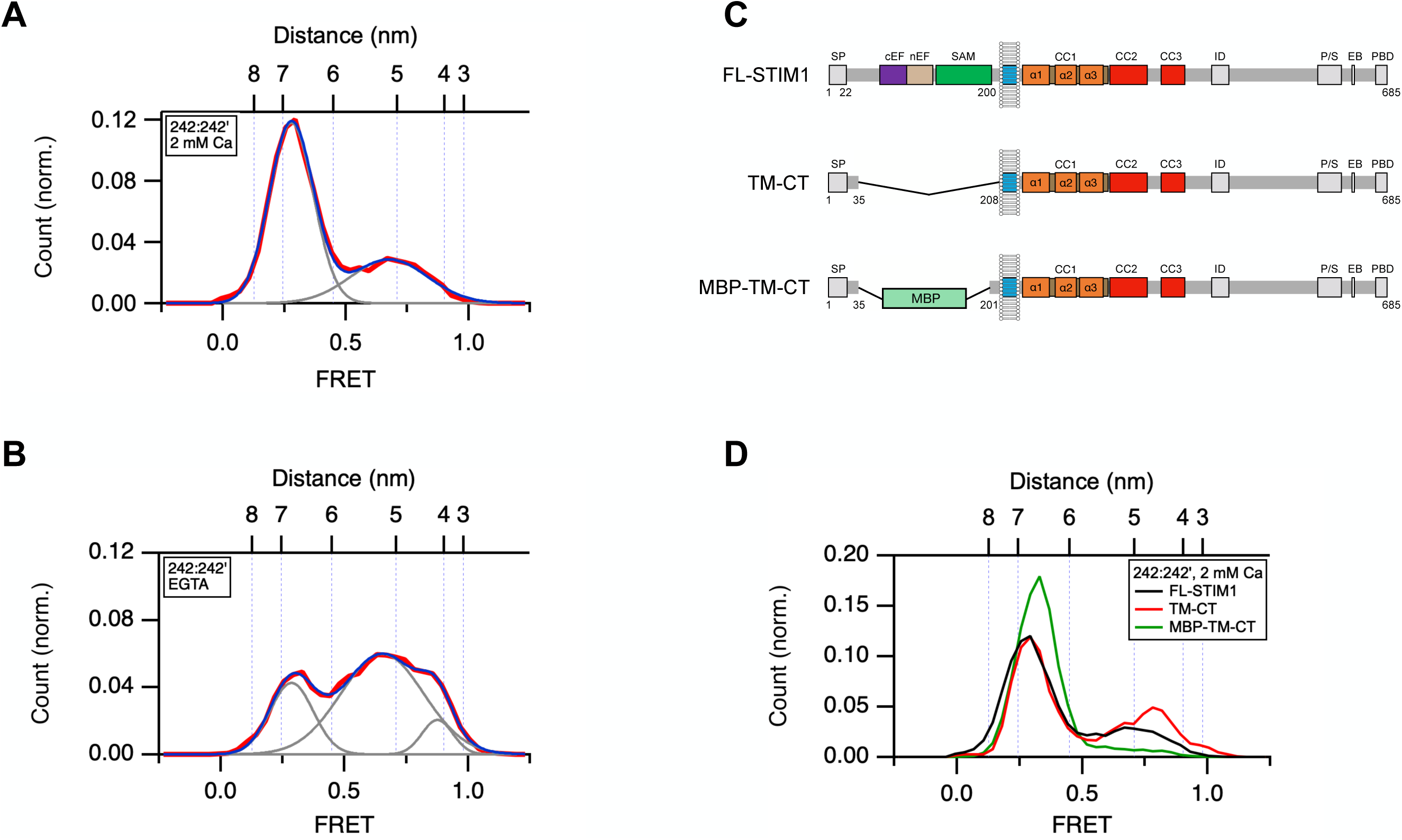
The Ca^2+^-bound EF-SAM domain inhibits spontaneous STIM1 activity through a steric mechanism. A, B. smFRET histograms for the 242:242’ dye pair in 2 mM Ca^2+^ (A; n=361) and 0.5 mM EGTA (B; n=437). Fitted Gaussian curves (gray) and their sum (blue) are superimposed on the data (red). The fractional area under the low-FRET (E=0.28) peak is used to indicate the probability of the resting conformation. C. Diagrams of full-length STIM1, TM-CT and MBP-TM-CT constructs. D. smFRET histograms for 242:242’ of FL-STIM1 (from panel A, n=361), STIM1 with deleted luminal domain (TM-CT; n=203), and STIM1 with MBP substituted for the luminal domain (MBP-TM-CT; n=234). Measurement of the fractional area of the low-FRET peaks is shown in **Supplementary** Fig. 4.

### The EF-SAM domain is a steric brake

Following store depletion, release of the EF hands from SAM is well recognized as the critical initiating step in STIM1 activation by allowing the SAM domains to dimerize (Sallinger et al., 2024; Stathopulos et al., 2008). A previous report that overexpression of an EF-SAM deletion mutant of STIM1 evoked spontaneous SOCE (Li et al., 2007) led us to hypothesize that the Ca^2+^-bound EF-SAM domain may also act as a steric brake on spontaneous activation. To test this using smFRET, we assessed changes in the resting state probability after removing the EF-SAM domain and after replacing it with maltose binding protein (MBP) (**Fig. 2C, D**). The probability of the resting state was measured from the area under the low-FRET peak from smFRET measurements of the 242:242’ dye pair. In the presence of saturating Ca^2+^, removal of the EF-SAM domain reduced occupancy of the low FRET state from 70% to 59% and increased the proportion of molecules with higher FRET, indicating destabilization of the resting state (**Supplementary** Fig. 4). Replacing the EF-SAM domain with maltose binding protein (MBP) reversed this effect and even enhanced the stability of the resting state beyond normal (91%). These results provide quantitative evidence that the Ca^2+^-bound EF-SAM domains create a brake on STIM1 activation by acting as a steric barrier to hinder the close apposition of the luminal and presumably the TM and CC1α1 domains, which form an extended coiled-coil in the active state (Hirve et al., 2018).

### Alignment of the CC1α1-CC3 interface positions the CAD apex at the ER membrane

The binding of CC1α1 to CC3 in CAD acts in two ways as a brake. In addition to keeping the CC1α1, TM, and luminal SAM domains well separated to reduce chances for spontaneous dimerization, it also distances the CAD apex, the region implicated in Orai1 activation (Wang et al., 2014; Butorac et al., 2019), as far as possible from the PM (**Fig. 1**). While multiple CC1α1 and CC3 residues that enforce the resting state have been identified through mutagenesis (Muik et al., 2009, 2011; Shrestha et al., 2022; Ma et al., 2015; Zhou et al., 2013), the registration of the interhelical interface is unknown. Both helices have hydrophobic faces which could potentially align in a number of ways, and their registration is important in part because it determines the position of the CAD apex relative to the ER membrane.

The AlphaFold2 model predicts close proximity of L258 with T420’, and of L261 with T420’ and L423’ (**Fig. 3A**). We tested these predictions using a cysteine-scanning approach, in which we introduced cysteines at various locations in mCh-STIM1 and co-expressed it in STIM1/2 DKO HEK293 cells with HA-STIM1 bearing the L258C or L261C mutation. The cells were then treated with the membrane-permeant oxidizing agent diamide, and the formation of disulfide-linked dimers was quantified by Western blot (see Methods) (**Fig. 3B, C; Supplementary** Fig. 5). Due to the domain-swapped interactions of CC1α1 and CC3’, close proximity of the introduced cysteines would enable formation of disulfide-linked heterodimers (**Fig. 1D**). L258C and L261C by themselves form some dimers after diamide treatment, which is consistent with ser, gly, or ala mutations at these sites weakening the CC1α1-CC3 clamp (Muik et al., 2011; Ma et al., 2015; Shrestha et al., 2022) and allowing transitions to the active CC1α1 coiled-coil state that enable disulfide bonds to accumulate over time. Quantitation of the disulfide heterodimer bands indicated that in the resting state, L258C is most proximal to T420C’ (**Fig. 3C**), while L261C is closest to T420C’ and L423C’ (**Supplementary** Fig. 5C). The formation of multiple disulfides is consistent with the AlphaFold2 structure and suggests flexibility in the side-chains. Together, these results support the ability of the AlphaFold2 model to describe the clamp interface and establish the alignment of the CC1α1 and CC3 helices in the resting state.

**Figure 3.**
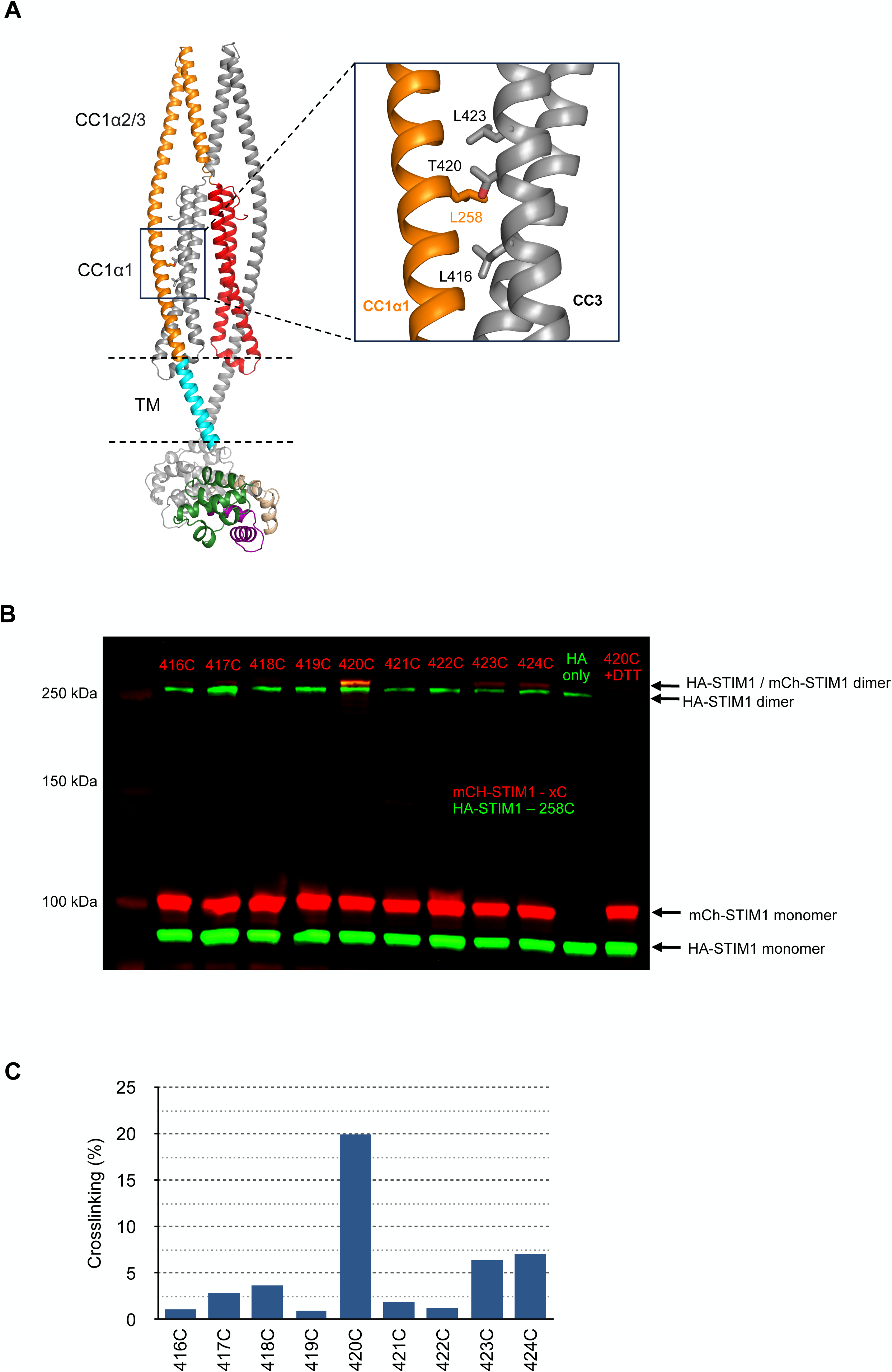
Helical alignment of the CC1α1-CC3 brake. A. Proximity of L258 (CC1α1) and T420 (CC3) predicted by the AlphaFold2 model. B. Western blot showing heterodimer disulfide crosslinking between HA-STIM1-L258C and mCh-STIM1-T420C. Each lane contains the lysate from diamide-treated cells coexpressing HA-STIM1-L258C and a single mCherry-STIM1 mutant (red) (see Methods). Treatment of the lysate with DTT removes the dimer bands, confirming disulfide crosslinking. Expression of HA-STIM1-L258C alone yields some homodimers (‘HA only’). C. Percent of heterodimers forming disulfide crosslinks, measured from the gel in B (see Methods). Results are representative of two experiments.

Interestingly, the registration between CC1α1 and CC3 predicts a close apposition of the CAD apex with the ER membrane, raising the possibility that protein-lipid interactions may influence the stability of the resting state. To address this, we examined the effects of lipid charge on the resting conformation probability in saturating Ca^2+^ as judged by smFRET of the 242:242’ dye pair. With liposomes made from 80% phosphatidylcholine (PC; no net charge) + 20% phosphatidylserine (PS; one net negative charge), probability of the low-FRET state was 71%, similar to the 70% observed with our standard lipid mixture having the same net charge [60% PC + 20% phosphatidylethanolamine (PE, uncharged) + 20% phosphatidylglycerol (PG, one net negative charge)] (**Fig. 4A, B**). In contrast, in uncharged liposomes containing only PC, the probability of the low-FRET state was reduced to 33% (**Fig. 4C**). These results demonstrate that negatively charged lipid headgroups act as a brake to stabilize the resting conformation of STIM1. A potential mechanism for this effect could involve the interaction of negatively charged headgroups with two clusters of basic residues (_382_KIKKKR) located near the apex of CAD (**Fig. 4D**). As a test of this idea, we mutated the four lysines to alanines (4KA) and repeated the smFRET measurements in standard liposomes (3 PC : 1 PE : 1 PG). The 4KA mutation reduced the probability of the resting conformation to 17% (**Fig. 4E**), supporting the notion that an electrostatic protein-lipid brake helps stabilize the resting state.

**Figure 4.**
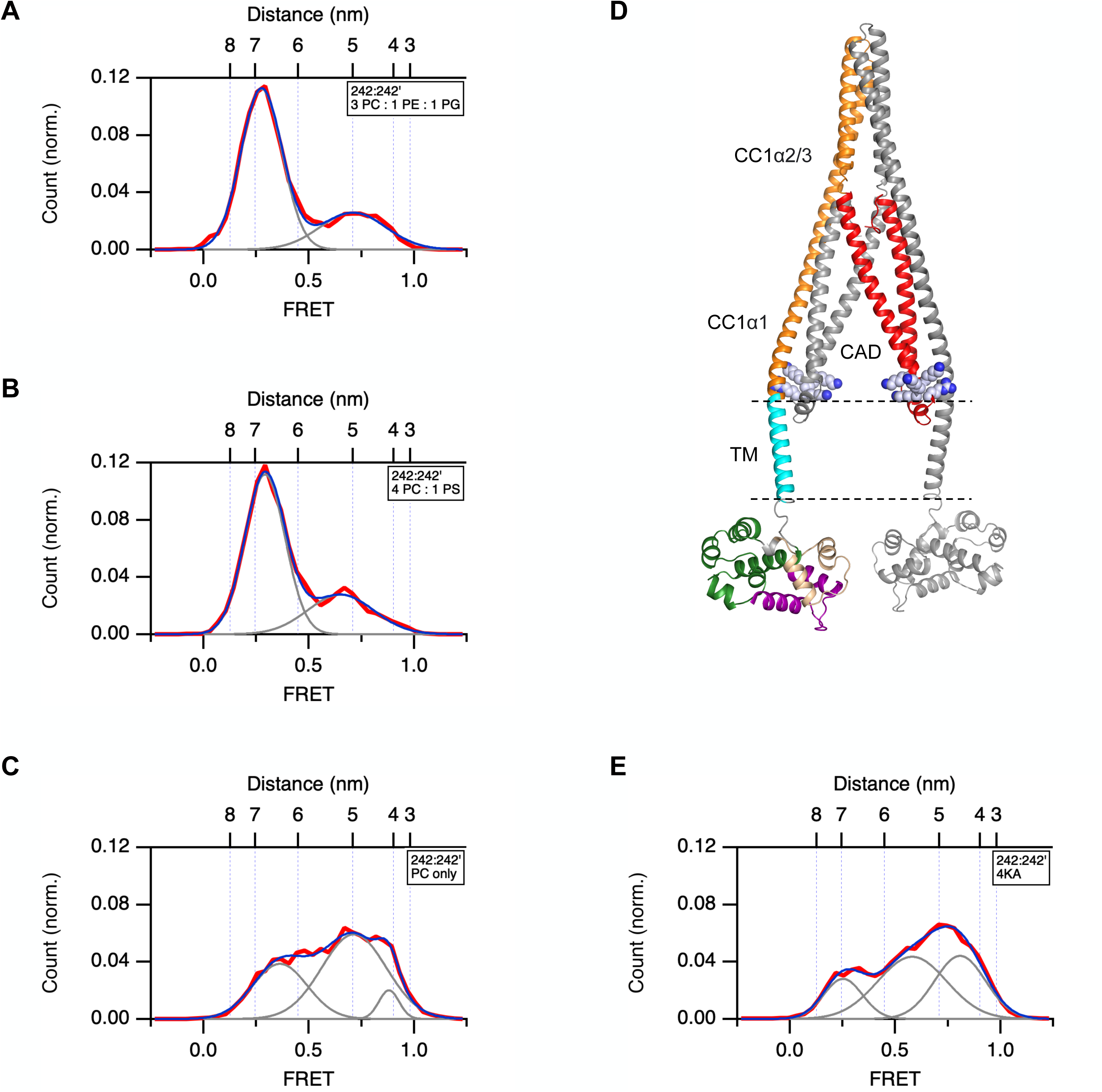
Electrostatic interaction of the CAD apex with the liposome membrane enhances the stability of the STIM1 resting state. In A-C, flSTIM1 was reconstituted in the presence of 2 mM Ca^2+^ in liposomes of varying composition and smFRET histograms were collected for 242:242’. Fitted Gaussian curves are superimposed to measure occupancy of the low-FRET state. A. Liposome composition 3 PC : 1 PE : 1 PG (20% negative charge; n=361). Histogram is reproduced from Fig. 2A. B. Liposome composition 4 PC : 1 PS (20% negative charge; n=204). C. Liposome composition PC only (uncharged; n=184). The lack of negatively charged lipids destabilizes the resting state. D. AlphaFold2 structure highlighting the position of the KIKKKR domain adjacent to the liposome membrane. Lysines and arginines are shown as spheres. E. STIM1-4KA (K382A/K384A/K385A/K386A) destabilizes the resting state when reconstituted in negatively charged liposomes containing 3 PC : 1 PE : 1 PG (20% negative charge; n=243).

### The CC1α2/3 dimer forms a fourth brake on STIM1 activity

Previous studies have shown that a quadruple mutation (E318A/E319A/E320A/E322A, or ‘4EA’) in CC1α3 partially activates STIM1 and SOCE (Korzeniowski et al., 2010; Yu et al., 2013; Muik et al., 2011; Zhou et al., 2013), suggesting a role for CC1α3 in stabilizing the resting state, but the underlying mechanism was unknown. We hypothesized that association of the CC1α2 and CC1α3 helices between the two subunits (**Fig. 1**) forms a brake on STIM1 activation, and that the 4EA mutation may activate STIM1 by destabilizing this structure. As shown in **Supplementary** Fig. 6A, 4EA reduces the occupancy of the high-FRET state for 309:309’ in the presence of Ca^2+^, indicating separation of the CC1α2/3 dimer. In addition, as shown by 242:242’ FRET, the 4EA mutation releases CC1α1 from CC3 in the presence of Ca^2+^ to a similar extent as EGTA does with WT STIM1 (cf. **Supplementary** Fig. 6B, **Fig. 2B**). Consistent with previous studies, STIM1-4EA evoked constitutive SOCE in unstimulated cells (**Supplementary** Fig. 6C).

To gain structural insight into the basis of CC1α2/3 as a brake, we used the AlphaFold2 model to identify residues that could stabilize the CC1α2/3 dimer in the resting state. The model predicts homotypic hydrophobic interactions between the two CC1α3 subunits at four locations: L321, V324, L328, and L335, as well as symmetric intersubunit salt bridges between CC1α2 and CC1α3’ at K294 and E332’ (**Fig. 5**). Mutating the four hydrophobic sites to serines (‘4S’ mutant) significantly destabilized the resting state as assayed by 242:242’ FRET (**Fig. 5B**). When expressed with Orai1 in HEK cells, the 4S mutant STIM1 evoked constitutive Ca^2+^ influx (**Fig. 5C**). The single mutations K294E or E332K produced an even stronger activating effect as judged by destabilization of the resting state in smFRET measurements (**Fig. 5E**) and stimulation of constitutive Ca^2+^ entry in cells (**Fig. 5F**). Importantly, the charge-swapped double mutant K294E/E332K returned STIM1 to its resting state as indicated by 242:242’ smFRET and restoration of normal resting [Ca^2+^]_i_ in cells (**Fig. 5E, F**), strongly supporting a direct interaction between the two sidechains. Together, these data reveal specific hydrophobic and electrostatic interactions that stabilize the CC1α2/3 dimer in the resting state to create a brake that minimizes spontaneous STIM1 activation.

**Figure 5.**
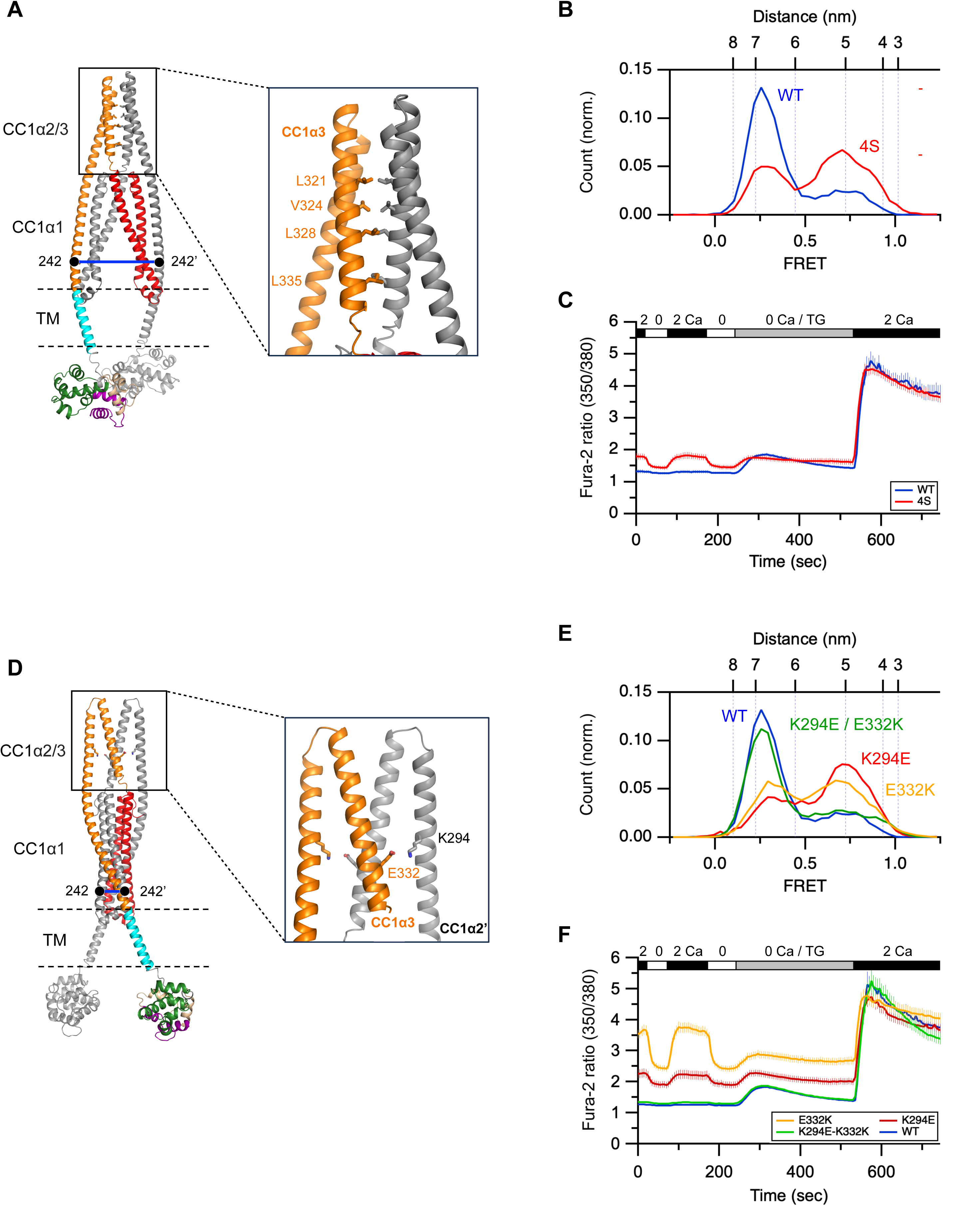
Structural basis of the CC1α2/3 brake. A. A symmetric hydrophobic interface between neighboring CC1α3 domains in the AlphaFold2 model. B. smFRET histogram of 242:242’ in STIM1-WT (from Fig. 2A) and STIM1-4S (L321S/V324S/L328S/L335S) in 2 mM Ca^2+^ (n=218). C. Fura-2 350/380 ratios (mean ± sem) from resting HEK cells expressing Orai1 and either STIM1-WT (n=57 cells) or STIM1-4S (n=63 cells). Changes in extracellular [Ca^2+^] (0 and 2 mM) and addition of thapsigargin (TG; 1 µM) are indicated. Resting [Ca^2+^]_i_ is elevated in cells expressing STIM1-4S, indicating constitutive activity of STIM1 and SOCE, while TG-induced SOCE is normal. D. Intersubunit salt bridges between K294 (CC1α2) and E332 (CC1α3) in the AlphaFold2 model. E. smFRET histogram of 242:242’ in STIM1-WT (from Fig. 2A) and STIM1-K294E (n=211), STIM1-E332K (n=216), or STIM1-K294E/E332K (n=255) in 2 mM Ca^2+^. F. Fura-2 350/380 ratios (mean ± sem) from resting HEK cells expressing Orai1 and either STIM1-WT (from C, n=57 cells), STIM1-K294E (n=62 cells), STIM1-E332K (n=87 cells), or STIM1-K294E/E332K (n=56 cells). Changes in extracellular [Ca^2+^] (0 and 2 mM) and addition of thapsigargin (TG; 1 µM) are indicated. Resting [Ca^2+^]_i_ is elevated in cells expressing STIM1-K294E or E332K, indicating constitutive activity of STIM1 and SOCE, but not in cells expressing the double mutant K294E/E332K. In all cases, TG-induced SOCE is normal.

### Membrane insertion of flSTIM1 affects conformational dynamics of the resting state

A comparison of smFRET values in flSTIM1 and ctSTIM1 (**Supplementary Table 2**) shows that membrane insertion of the TM domains in flSTIM1 does not affect the predominant structure of the cytosolic domain; however, it does create significant differences in conformational dynamics. In ctSTIM1, we previously observed frequent switching between domain-swapped and non-swapped configurations of the CC1α1-CC3 clamp (van Dorp et al., 2021). To compare clamp switching in ctSTIM1 and flSTIM1, we counted FRET transitions for the 242:431’ dye pair occurring within the first 5 s of single-molecule traces. While ctSTIM1 displayed frequent transitions, flSTIM1 only rarely fluctuated, with molecules showing constant FRET values for up to tens of seconds (**Fig. 6**). These results show that membrane insertion of flSTIM1 helps to stabilize the domain-swapped configuration of the CC1α1-CC3 clamp.

**Figure 6.**
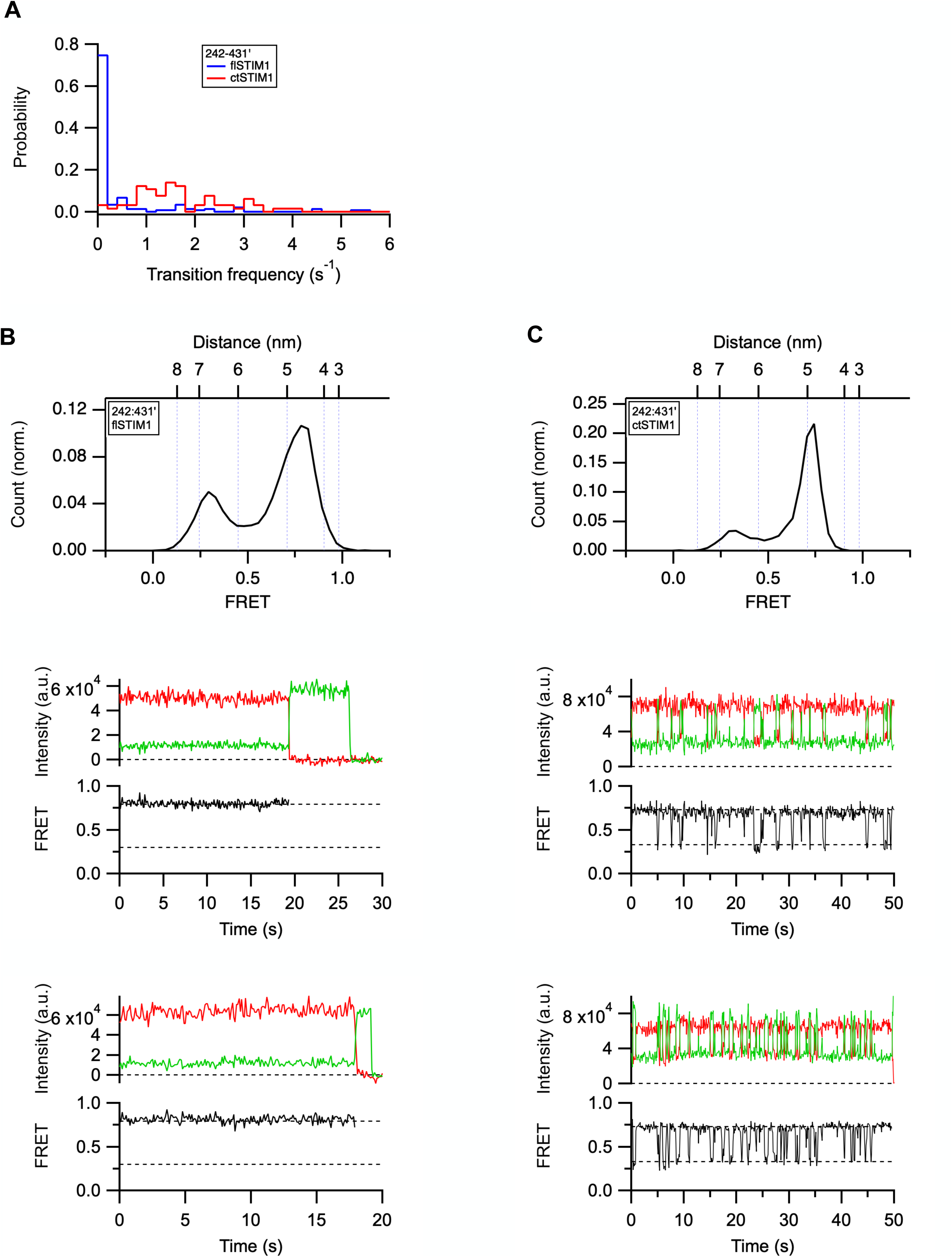
Membrane insertion of flSTIM1 reduces conformational fluctuations out of the resting state. A. Probability distribution of the mean smFRET transition frequency over a 5-s period for flSTIM1 and ctSTIM1, measured for the 242:431’ dye pair (CC1α1-CC3); n=189 (flSTIM1) and n=78 (ctSTIM1). B. smFRET histogram for flSTIM1 242:431’ (top; from Fig. 1C), and representative traces (bottom) of donor (green) and acceptor (red) fluorescence and FRET (black). Acceptor photobleaching followed by donor photobleaching is shown. Dashed lines indicate peak values from the smFRET histogram. Note the long stable occupancy of the high-FRET resting state. C. smFRET histogram for ctSTIM1 242:431’ (top), and representative traces (bottom) of donor (green) and acceptor (red) fluorescence and FRET (black). Dashed lines indicate peak values from the smFRET histogram. ctSTIM1 makes frequent transitions out of the high-FRET resting state due to switching between domain-swapped and non-swapped conformations (see text).

## DISCUSSION

In this study we used smFRET and AlphaFold2 to describe the conformation of full-length STIM1 in a membrane environment in the Ca^2+^-bound state, and to identify the structural restraints that maintain it. We present evidence that the stability of the inactive Ca^2+^-bound state is determined by at least four weak brakes, all of which are required to minimize spontaneous activation of STIM1.

### A description of the resting state

The inactive structure of flSTIM1 in membrane has several key features based on distances derived from smFRET measurements. CC1α1 is closely associated with CC3 of CAD in a parallel orientation, positioning the apex of CAD near the membrane. This configuration is an effective way of distancing the apex, which is thought to interact with Orai1 to open the channel (Wang et al., 2014; Calloway et al., 2010; Korzeniowski et al., 2010; Thompson et al., 2018; Butorac et al., 2019), as far as possible from the plasma membrane when ER Ca^2+^ stores are full. The interaction is domain-swapped, with each CC1α1 helix interacting with the CC3 of the neighboring subunit. By providing additional connections between the two STIM1 subunits, domain-swapping further stabilizes the dimeric structure beyond what is provided solely by interactions of the two CC2-CC3 hairpins within the basal region of CAD (Yang et al., 2012).

Finally, the CC1α2 and CC1α3 helices form a dimer of hairpins that point away from the base of CAD. These results clarify for the first time the overall arrangement of helices and domains in the resting state of full-length STIM1 embedded in membrane.

These features of the inactive resting state are similar to those we described in a previous smFRET study of the soluble cytosolic fragment ctSTIM1 (van Dorp et al., 2021), showing that they are not dependent on membrane insertion of the TM domains (**Supplementary Table 2**). However, the conformational dynamics of flSTIM1 differ significantly from those of ctSTIM1. Compared to ctSTIM1, in which the CC1α1-CC3 clamp transitions frequently between domain-swapped and non-swapped conformations, the clamp in flSTIM1 is much more stable, with many molecules lacking fluctuations for up to tens of seconds (**Fig. 6**). This increased stability may result from membrane insertion as well as electrostatic interaction of lipid headgroups with the CAD apex (**Fig. 4**), and it highlights the advantages of membrane-reconstituted flSTIM1 as a model system for identifying the physical interactions that govern STIM1 activity.

We were surprised to find that the luminal domains in flSTIM1 are quite dynamic, even in the presence of saturating (2 mM) Ca^2+^. In the NMR structure of Ca^2+^-bound EF-SAM, the cEF and nEF hands are wrapped around helix 10 of the SAM domain, effectively preventing SAM-SAM dimerization and STIM1 activation (Stathopulos et al., 2008). The association between the cEF hand and SAM is not entirely stable even in saturating Ca^2+^. Using the 64:178 dye pair as a probe, the FRET histogram indicates multiple states, and transitions out of the predominant high-FRET state of 0.9 suggests transient release of the cEF hand from SAM (**Supplementary** Fig. 7A). Thus, even in the presence of saturating Ca^2+^ the SAM domain may intermittently become available for dimerization. In fact, the SAM-SAM distance measured with a 178:178’ dye pair also fluctuated over a wide range (**Supplementary** Fig. 7B), suggesting that even under Ca^2+^-saturated resting conditions, they may collide. Thus, the transient release of SAM from the cEF hand coupled with random collisions between SAM domains may allow luminal domains to dimerize, setting limits on the basal activity of STIM1 in resting cells.

We selected an AlphaFold2 model for the flSTIM1 dimer that was consistent with smFRET-derived distances throughout the cytosolic domain (**Fig. 1**). The structure proved to be accurate enough to reveal interactions of sidechains in critical regions that control STIM1 activity. The model correctly predicted CC1α1-CC3’ interaction sites that were confirmed through cysteine scanning (**Fig. 3, Supplementary** Fig. 5), as well as electrostatic interactions between lysine clusters in the CAD apex and the liposome membrane (**Fig. 4**), and intersubunit hydrophobic and electrostatic interactions that stabilize the CC1α2/3 dimer in the resting state (**Fig. 5**). The model can benefit from further refinement; for example, it predicts a continuous helix from the TM domain through CC1α2, despite the likelihood of non-helical linkers at the border between TM and CC1α1 (aa 233-236) and between CC1α1 and CC1α2 (aa 273-275), based on bioinformatics analysis (Soboloff et al., 2012) and an NMR structure of the CC1 monomer (Rathner et al., 2020). The absence of these flexible linkages in the AlphaFold2 model may alter the rotation of helices relative to each other, which may explain why the TM 222:222’ distance in the model was not consistent with smFRET data (see Methods).

### Multiple weak brakes control STIM1 activation

In addition to describing the structure of Ca^2+^-saturated STIM1 in the inactive state, our AlphaFold2 model and additional smFRET measurements reveal the structural basis for four brakes that work together to minimize spontaneous activation: the EF-SAM domain, the CC1α1-CC3 clamp, the CAD apex-lipid interaction, and the CC1α2/3 dimer.

*EF-SAM domain.* Several reports have described spontaneous puncta formation and SOCE in cells overexpressing a truncated STIM1 variant lacking the EF-SAM domain, suggesting that the Ca^2+^-bound luminal domain itself may suppress spontaneous activity (Li et al., 2007; Muik et al., 2011; Ma et al., 2017). Our results confirm these findings and suggest a steric mechanism for this effect. Removal of the luminal domain caused a modest degree of activation detected by smFRET, and this was prevented by luminal addition of MBP, a protein comparable to but slightly larger than the EF-SAM domain. Based on these results, we suggest that the bulk of the Ca^2+^-bound EF-SAM domains in STIM1 prevents close approach of the TM and CC1α1 domains, thereby limiting the opportunity for dimerization of the TM and CC1 domains and spontaneous activation. It is important to recognize that activation caused by deletion of the EF-SAM domain is quite modest compared to that produced by Ca^2+^ removal from the WT protein (cf. **Fig. 2B, D**), highlighting the importance of SAM-SAM dimerization for stabilizing and promoting the active state. Thus, the luminal domain performs two opposite functions: in the Ca^2+^-bound state it acts as a brake to help keep STIM1 inactive in resting cells, while in the Ca^2+^-free state it becomes an ‘accelerator’ to promote STIM1 activation through SAM-SAM dimerization.

*CC1α1-CC3 clamp.* Pioneering work by Romanin, Hogan, Zhou and their colleagues established the importance of a CC1α1-CC3 interaction in limiting spontaneous activation of STIM1 (Fahrner et al., 2014; Ma et al., 2015, 2020; Shrestha et al., 2022; Muik et al., 2009, 2011). Our structural model reveals several mechanisms by which the CC1α1-CC3 clamp acts to inhibit activation. First, CC1α1 binding to CC3 keeps both the TMs and the luminal domains well separated in the resting state (**Fig. 1**), limiting the opportunities for interaction that could lead to spontaneous activation. The large distance between TM domains indicated by smFRET contrasts with a previous proposal that the TMs are associated in the resting state, and in which activation occurs through rotation of the crossed helices to allow formation of a coiled-coil (Ma et al., 2015; Korzeniowski et al., 2017). We believe instead that activation must involve a significant translational movement of the separated luminal domains to bring SAM domains into contact for dimerization. The discrepancy with our results likely arose from the use of TM-CC1 fragments in the previous study, which can bind weakly to each other in the absence of the V-shaped CAD (Covington et al., 2010), and possibly from the use of a TM mutation (C227W) rather than Ca^2+^ depletion to drive activation, which may have biased the interaction of the TM domains.

A second essential aspect of the CC1α1-CC3’ brake is that the parallel orientation of the two helices orients the CAD apex toward the ER membrane, distancing it as far as possible from the plasma membrane and Orai1. In addition to their parallel orientation, the register between the helices is a key factor that determines the proximity of the CAD apex to the ER membrane in the resting state. Several different computational models for the registration between CC1α1 and CC3 have been published based on hypothetical alignments of residues known by mutagenesis to be required for the clamp. These include antiparallel CC1α1 and CC3 helices with L258 paired with V419 (Ma et al., 2015), as well as parallel helices with L258 paired with L423 (van Dorp et al., 2021) or with L416 (Horvath et al., 2023). To experimentally determine the register, we adopted a cysteine scanning approach that was previously used to identify the pore-lining residues of Orai1 (McNally et al., 2009; Zhou et al., 2010) and the locations of interacting CC1 sidechains in activated STIM1 (Hirve et al., 2018; van Dorp et al., 2021). Disulfide formation revealed proximity of residues L258 to T420’ and of L261 to T420’ and L423’, as predicted by our AlphaFold2 model (**Fig. 3**, **Supplementary** Fig. 5). This result differs from all previous models; it places CAD one helical turn further from the membrane than van Dorp et al (van Dorp et al., 2021), one turn closer to the membrane than Horvath et al (Horvath et al., 2023), and several turns closer than Höglinger et al (Höglinger et al., 2021). Interestingly, the AlphaFold2 model also predicts interactions of L248 with I409’, perhaps explaining why alanine substitutions at either site release CAD in a split STIM1 assay (Shrestha et al., 2022). In addition to describing the interaction interface in the CC1α1-CC3 clamp, the model predicts that the CAD apex may extend beyond the edge of the membrane (as inferred by the position of Q233). Membrane insertion could help explain why large hydrophobic residues at position 394 in the apex (phe, leu) appear to stabilize the inactive state (Höglinger et al., 2021; Zhou et al., 2023), although other mechanisms are possible, and insertion may not actually occur due to the flexibility of the apex (van Dorp et al., 2021).

*CAD apex-lipid interaction.* Regardless of whether the CAD apex inserts into the membrane, the registration of the CC1α1-CC3 clamp positions it close enough to interact with lipid headgroups. Neutralizing either the membrane charge (PC liposomes) or the two KIKKKR regions (4KA) destabilized the resting state, suggesting the operation of an electrostatic brake (**Fig. 4**). An electrostatic interaction between the ER membrane and the 10 positive charges of the KIKKKR domains in cells would be expected given the composition of the ER membrane, which contains ∼17% negatively charged lipid (PS and phosphatidylinositol) (Van Meer et al., 2008), and physical modeling studies predicting a −25 mV isopotential electric field at ∼1 nm from a 2:1 PC/PS membrane (Murray et al., 2002). Interestingly, in store-depleted cells where the CAD is extended away from the ER, the KIKKKR region has been reported to interact with negatively charged PI4P in the PM to help trap STIM1 at ER-PM junctions (Cohen et al., 2023) and to enhance binding to Orai1 (Calloway et al., 2010; Korzeniowski et al., 2010). Thus, like the EF-SAM domain and the CC1 domain, the KIKKKR domain appears to serve two opposing functions: to stabilize the inactive state under resting conditions and to promote the activated state after store depletion.

*CC1α2/3 dimer.* A role for CC1α3 in stabilizing the STIM1 resting state was first proposed by Balla and colleagues, who reported that the 4EA mutation (E318A/E319A/E320A/E322A) in

CC1α3 caused partial activation of STIM1 and SOCE (Korzeniowski et al., 2010). Later studies showed that the 4EA mutation releases CC1α1 from CAD in ctSTIM1 (Zhou et al., 2013) and causes a small extension of the OASF STIM1 fragment (aa 233-474) consistent with activation (Muik et al., 2011). Several hypotheses have been proposed to explain how the EEELE domain stabilizes the resting state. Initially, 4EA was thought to disrupt an electrostatic interaction of the glutamates with the basic KIKKK residues near the CAD apex (Korzeniowski et al., 2010), but later studies ascribed the activating effects instead to disruption of amphiphilic helical properties (Yu et al., 2013) or stabilization of a potential CC1α3 binding site for Orai1 (Stathopulos et al., 2013). Ma et al suggested that CC1α3 may be important for maintaining CC1α1 and CC3 in a position to interact (Ma et al., 2015). Our results are compatible with this idea and suggest an underlying mechanism by which the amphiphilic CC1α3 helix allows the CC1α2/3 dimer to form, which in turn enhances CC1α1 binding to CC3.

To describe a structural basis for the CC1α2/3 brake, we identified potential interaction sites between the two CC1α2/3 hairpin structures in the AlphaFold2 model that would be expected to stabilize the resting state structure in which the two hairpins are closely connected. A hydrophobic interface between CC1α3 domains and an intersubunit electrostatic interaction between CC1α2 and CC1α3 were identified, and mutations at these sites caused varying degrees of spontaneous STIM1 activation as assayed by smFRET for the 242:242’ (CC1α1) dye pair and constitutive SOCE in resting cells. The activating effects of charge reversal at either K294 or E332 give strong evidence for an electrostatic interaction that holds the two CC1α2/3 hairpins together, which is further strengthened by the normal behavior of the charge-swapped double mutant K294E/E332K. These results outline the structural basis for a new regulatory domain of STIM1, the CC1α2/3 dimer, in which the interacting pair of CC1α2/3 hairpin domains stabilizes the interaction of CC1α1 with CAD. We propose that it does this by holding the CAD in position, so that when CC1α1-CC3 contacts are transiently broken, the CAD preferentially rebinds to CC1α1 rather than escape. This mechanism can explain why in the split STIM1 assay, in which the CC1α3 is not connected directly to CC2 of CAD and therefore cannot hold CAD in position, the CC1α2 and CC1α3 domains fail to enhance CAD trapping beyond what is achieved by the CC1α1 domain alone (Ma et al., 2015).

### Implications of multiple weak brakes for the regulation of STIM1 and SOCE

The association of GOF and LOF mutations in STIM1 and Orai1 with human disease shows that to avoid pathological consequences, STIM1 needs to be strongly suppressed when ER Ca^2+^ stores are replete, yet readily activated by receptor-triggered loss of ER Ca^2+^, and reliably deactivated upon store refilling. The question is how these potentially conflicting requirements can be met. The simplest regulatory mechanism would involve a single strong brake that keeps STIM1 inactive under Ca^2+^ saturated conditions; in this case, the release of Ca^2+^ upon store depletion would enable dimerization of SAM domains to overcome the brake and drive the subsequent conformational changes (TM-CC1 coiled-coil formation, release of the PBD and CAD for binding to the PM and Orai1) that lead to Ca^2+^ entry. In order for such a single brake to reliably suppress STIM1 in the face of spontaneous fluctuations in the luminal domain (**Supplementary** Fig. 7), it would have to be strong. To overcome such a strong brake and enable robust activation, the SAM-SAM binding would need to be tight, which would make reversal of activation upon Ca^2+^ refilling energetically unfavorable.

Here we have shown the operation of at least four brakes that restrain the activation of STIM1 under resting, Ca^2+^-saturated conditions. The activating effects of mutations show that these brakes are all relatively weak. Releasing any one brake by mutating interface residues, e.g., L248S or L251S for the CC1α1-CC3’ clamp (Muik et al., 2011), KIKKK>AIAAA for the CAD apex-lipid brake (**Fig. 4**), or K294E or E332K for the CC1α2/3 brake (**Fig. 5**), is enough to unmask significant spontaneous activation, demonstrating that in each case the remaining three brakes together are sufficient to maintain the inactive state. In this case, the total energetic stabilization afforded by four brakes in tandem can resist spontaneous activation, and SAM dimerization now only needs to supply enough energy to overcome one of the brakes, e.g., the CC1α1-CC3’ clamp, because the remaining brakes are too weak to prevent further activation. A moderately weak SAM-SAM interaction ensures that upon store refilling, the Ca^2+^-bound cEF hand will have a chance to rebind to SAM and return STIM1 to its resting state. In this way, STIM1 can be reliably inactive when stores are full yet reversibly activated by store depletion, as required for physiological control of Ca^2+^ entry through store-operated Orai1 channels. These studies are a first step in understanding the full structural basis for STIM1 stability and reveal multiple intersubunit interaction sites that may ultimately provide pharmacological targets to modulate STIM1 and SOCE in vivo.

## METHODS

### Cell culture

HEK 293 cells were cultured in DMEM containing 2 mM L-alanyl-glutamine, 10% FBS, and 100 U/ml penicillin/streptomycin at 37°C in 5% CO_2_. Sf9 cells were cultured in ESF 921 insect cell culture medium (Expression Systems) at 27°C with constant shaking at 120 RPM. Baculovirus was generated with Bac-to-Bac baculovirus expression system (Invitrogen).

### DNA constructs

For insect cell expression, STIM1 constructs were cloned in the pFastBac1 vector by PCR from full-length human STIM1 (Origene) as follows. To construct his_6_-STIM1, STIM1 residues 1-34 followed by a C-terminal his_6_-tag and 3C protease cleavage sequence (LEVLFQGP) were inserted between EcoRI and XhoI restriction sites, and STIM1 residues 35-685 were inserted between XhoI and KpnI restriction sites. To construct MBP-STIM1, STIM1 residues 1-34 were inserted between BamHI and EcoRI restriction sites, the MBP sequence with a C-terminal 3C protease cleavage sequence was inserted between EcoRI and XhoI restriction sites, and STIM1 residues 35-685 were inserted between XhoI and KpnI restriction sites.

All native cysteines in STIM1 (C49, C56, C227 and C437) were mutated to serines, and desired mutations were introduced by site-directed mutagenesis (Quikchange XL; Stratagene). The activity of the cysteineless flSTIM1 appeared to be normal as assessed by resting [Ca^2+^]_i_ and thapsigargin-induced SOCE in HEK 293 cells cotransfected with Orai1 (**Supplementary** Fig. 1). This result is consistent with normal puncta formation by a C227S/C437S STIM1 mutant after TG treatment (Hirve et al., 2018) but differs from another study in which a single C437S mutation was reported to inhibit STIM1 activation (Kodakandla et al., 2022). The reasons for this discrepancy are unclear.

His_6_-TM-CT was made by replacing residues 35-685 of his_6_-STIM1 with STIM1 residues 208-685. His_6_-MBP-TM-CT was made by first linking MBP and 201-685 of STIM1 using Gibson assembly kit (New England Biolabs, E2611L) and substituting for residues 35-685 of his_6_-STIM1.

For Ca^2+^ imaging in HEK 293 cells (**Fig. 5, Supplementary** Fig. 6), the cysteineless STIM1 and other STIM1 mutants were generated from mCherry-STIM1 (Wu et al., 2006) by site-directed mutagenesis (Quikchange XL; Stratagene). For cysteine crosslinking experiments in HEK 293 cells (**Fig. 3, Supplementary** Fig. 5), constructs were generated from mCherry-STIM1 and HA-STIM1 (Wu et al., 2006) by site-directed mutagenesis (Quikchange XL; Stratagene). C437, the only cysteine in the cytosolic domain, was mutated to serine and the desired residues were mutated to cysteine for diamide-induced crosslinking.

All plasmids were verified by Sanger sequencing or whole plasmid sequencing.

### Protein expression and purification

All STIM1 constructs in pFastBac1 vector were expressed in Sf9 cells infected with recombinant baculovirus (Bac-to-Bac, Invitrogen). For symmetric inter-subunit smFRET samples, baculovirus of His_6_-tagged STIM1 with a single cysteine substitution was used for expression. For asymmetric inter-subunit and intra-subunit smFRET samples, baculoviruses of His_6_-tagged STIM1 and MBP-tagged STIM1 were used together for heterodimer expression.

Sf9 cells were harvested 45 h post infection and were lysed in a buffer of 20 mM Tris pH 7.8, 0.5 mM EDTA, 500 µM TCEP with added protease inhibitor cocktail (Sigma-Aldrich, S8830). Cell membranes were centrifuged at 150,000 × g for 30 min at 4°C. STIM1 protein was extracted from the membranes using a buffer of 20 mM Tris pH 7.2, 150 mM NaCl, 2% n-dodecyl-β-D-maltoside (DDM), 10 mM imidazole, 500 µM tris(2-carboxyethyl)phosphine (TCEP) and benzonase (1 µl/100 ml; Sigma-Aldrich) for 1 h at 4°C. After centrifugation, Ni-NTA resin was added to the supernatant and rotated for 2 h at 4°C. Ni-NTA resin was washed with 20x column volume of wash buffer containing 20 mM Tris pH 7.2, 150 mM NaCl, 0.1% DDM, 500 µM TCEP, and 40 mM imidazole. Protein was eluted in wash buffer supplemented with 300 mM imidazole. To isolate heterodimers, amylose resin was added to the elution and rotated for 1 h at 4°C, and protein was eluted in 20 mM Tris pH 7.2, 150 mM NaCl, 0.1% DDM, 500 µM TCEP, and 10 mM maltose. To cleave the tag, 3C protease was added to the eluted protein for 1 h at room temperature. Complete cleavage was verified by SDS-PAGE. After concentrating with a 100 kDa cutoff concentrator (Amicon Ultra), the STIM1 protein was further purified by size exclusion chromatography (SEC) using a Superose 6 Increase 10/300 GL column (Cytiva) equilibrated with SEC buffer of 20 mM Tris pH 7.2, 150 mM NaCl, 0.1% DDM, and 100 µM TCEP. Fractions containing STIM1 protein were collected and concentrated with a 100 kDa cutoff concentrator (Amicon Ultra). The STIM1 protein was supplemented with 10% (v/v) glycerol and flash frozen in liquid nitrogen. The purity of STIM1 samples were higher than 90% as determined by SDS-PAGE.

### Protein labeling and reconstitution in liposomes

Sites for cysteine substitution and dye labeling were selected from outward-facing residues if a structure was available (EF-SAM, CAD), and if not, from residues predicted to lie outside interface surfaces. STIM1 dimer containing two cysteines was labeled with maleimide-conjugated Alexa Fluor 555 and Alexa Fluor 647 (ThermoFisher). STIM1 was diluted to 10 µM in 200 µl desalting buffer (20 mM Tris pH 7.2, 150 mM NaCl, 0.1% DDM, and 100 µM TCEP). 5 µM donor fluorophore and 5 µM acceptor fluorophore were added and after incubation for 1 h at 4°C, free dye was removed with a home-packed desalting column filled with 5 ml of G50 resin (Sigma) pre-equilibrated with desalting buffer. Labeling efficiency was ∼50% for most samples. The labeled STIM1 protein was concentrated to ∼10 µM, aliquoted, supplemented with 10% (v/v) glycerol and flash frozen.

To reconstitute STIM1 proteins in liposomes, PC, PE, and PG (Avanti) were mixed at a 3:1:1 ratio in chloroform, and 0.5% biotinylated PE (Avanti) was added. The lipids in chloroform were dried by nitrogen gas and kept in vacuum overnight. The dry lipids were resuspended in buffer containing 20 mM Tris pH 7.2 and 150 mM NaCl to a final concentration of 10 mg/ml, creating a cloudy solution. Using an extruder set (Avanti Polar Lipids) the lipids were then extruded through a membrane with 100-nm pores (Whatman) 30 times to form liposomes with a diameter of ∼100 nm. For each sample, 180 µl of liposomes (10 mg/ml) was added to 20 µl of 12% n-octyl-β-D-glucopyranoside (βOG) solubilized in the same buffer and rotated for 15 min at 4°C. 10 µl of labeled STIM1 protein (∼10 µM) was added to the liposomes and rotated for 30 min at 4°C. 25 mg of Bio-Beads (Bio-Rad) was added to the sample and rotated for 1 h at 4°C, and another 25 mg of Bio-Beads was added and rotated for 1 additional hour. The reconstituted proteoliposomes were separated from free detergent and free STIM1 protein using a home-packed Sepharose CL-4B column (Sigma Aldrich). The fraction containing STIM1 proteoliposomes was added to 10 mg Bio-Beads and rotated overnight at 4°C to remove residual detergent, at which point the sample was ready for smFRET experiments.

### Imaging chamber preparation

Imaging chambers for single-molecule fluorescence imaging were prepared according to established protocols (Roy et al., 2008). Briefly, microscope slides and coverslips were first cleaned by stepwise sonication for 15 min each in glass containers with acetone, ethanol, 1 M KOH and pure water. They were then coated with a 100:1 PEG/PEG-biotin mixture (Laysan Bio) prior to flow cell construction. Strips of double-sided tape were applied to a quartz microscope slide (Finkenbeiner) to form channel walls, with holes at both ends of each channel to allow exchange of sample solutions. A microscope coverslip of 1.5 thickness (Erie Scientific) was pressed on the tape strips and edges of the channels were sealed with epoxy glue (Devcon).

Prior to attaching biotinylated liposomes to the coverslip surface, channels were rinsed with 20/150 TBS (150 mM NaCl, 20 mM Tris, pH 7.2 with HCl) then incubated with 0.2 mg/ml neutravidin (Thermo Fisher CWA) for 5 min. Liposomes were loaded into the channels after washing out the neutravidin. Prior to TIRF imaging, channels were filled with 20/150 TBS containing 100 μM cyclooctatetraene (Sigma Aldrich) and an oxygen scavenging system consisting of 1 % D-glucose, 1 mg/ml glucose oxidase (Sigma Aldrich), and 0.04 mg/ml catalase (Sigma Aldrich).

### TIRF microscopy and smFRET measurements

All smFRET experiments were performed at room temperature following a previous protocol (van Dorp et al., 2021) with some modifications. The home-built TIRF system is based on an Axiovert S100 TV microscope equipped with a Fluar 100x 1.45 NA oil-immersion objective (Zeiss). 532- and 637-nm lasers (OBIS 532 nm LS 150 mW, Coherent and OBIS 637 nm LX 140 mW, Coherent) were used for excitation in objective TIRF mode. Donor and acceptor signals were separated by a 652 nm dichroic (Semrock) and passed through 580/60 nm and 731/137 nm bandpass filters (Semrock), respectively, mounted in an OptoSplit-II beamsplitter (Cairn Research) to an EM-CCD camera (iXon DU897E, Andor). Hardware and data acquisition were controlled by homemade scripts in μManager.

For each sample, the molecule density and distribution on the coverslip surface was checked with the TIRF microscope. When optimal density was achieved (300 to 500 molecules in the camera field of view), donor and acceptor emission were recorded with a 100-ms integration time under 532-nm laser excitation for 80 s, immediately followed by 1 s of 637-nm laser excitation to verify acceptor dye bleaching.

Data were analyzed using custom Python scripts as detailed previously (van Dorp et al., 2021). Briefly, molecules with single acceptor bleaching before single donor bleaching were selected. The FRET ratio E was calculated at each time point as E = I_A_/(I_A_ + γI_D_), where I_A_ and I_D_ are the acceptor and donor fluorescence values, respectively, and γ was measured empirically for each molecule as described previously (van Dorp et al., 2021). For single-molecule traces longer than 20 points (2 s), summed FRET histograms were constructed by distributing FRET amplitudes into 40 bins (from –0.25 to 1.25) and normalizing by the number of points in each trace. Histograms were fitted with a sum of Gaussian functions using routines written in Igor Pro (WaveMetrics).

R_0_ of the dye pair was measured empirically on our smFRET system. The crystal structure of CAD was used to simulate the dye-dye distances for 417-417’ and 431-431’ pairs using Crystallography and NMR System (CNS) (Brunger, 2007). These distances were used to solve for R_0_ using the FRET equation and 417:417’ and 431:431’ smFRET measurements from ctSTIM1 samples. The calculated R_0_ values were 5.85 nm for 417:417’ and 5.80 nm for 431:431’; a value of 5.8 nm was used to estimate distances throughout this paper. Distances (R) were calculated from FRET (E) using the relation E = 1/[1 + (R/R_0_)^6^].

### Dye position simulation with Crystallography and NMR System (CNS)

We simulated dye positions on STIM1 structures using Crystallography and NMR System (CNS) as described (Choi et al., 2010). Briefly, the pdb file was loaded in Pymol and residues for dye labeling were mutated to cysteine. Atomic models of the fluorophores and their maleimide linker were attached to the mutated cysteines and subjected to molecular dynamics simulations by fixing all protein atoms except the dye and linker atoms. 100 simulations were performed for each labeling pair and the average distance between the dye centers (CAO atom) was calculated from the resulting coordinates.

### Generating structural models with AlphaFold2

Residues 35-444 of human STIM1 was used to generate AlphaFold2 models, with the following parameters: Num_relax = 0; Template_mode = none; MSA mode = mmseqs2_uniref_env; Pair_mode = unpaired_paired; Model_type = auto; Num_recycles = 48; Recycle_early_stop_tolerance = auto; Relax_max_iterations = 200; Pairing_strategy = greedy. Out of the top five models shown in **Supplementary** Fig. 3, only model 2 shows domain-swap binding between CC1 and CAD as well as overall agreement with smFRET-derived distances.

In all structures, only residues from the cEF hand to the N-terminal end of CAD (63-444) are shown. Residues beyond this location are predicted to be unstructured and showed low confidence in the AlphaFold2 model (pIDDT values <0.5). While the model accurately predicted inter-dye distances estimated from smFRET in the cytosolic region, it needs refinement for the TM and luminal domains. The model underestimated the TM spacing (222:222’; smFRET distance 6.4 nm, model distance 4.1 nm), which may be related to the absence of a flexible linker predicted by JPRED3 to connect the TM domain to CC1α1 (Soboloff et al., 2012), possibly resulting in a rotation of the TM helices that orients the 222 residue sidechains to face each other. In addition, while the cEF hand-SAM distance (64:178) from smFRET (4.1 nm) is consistent with dye positions added to the published EF-SAM NMR structure (4.1 nm), it differs from the AlphaFold2 model (2.0 nm). For these reasons, we only applied the model to make predictions about the cytosolic region.

### Ca^2+^ imaging

HEK293 cells were co-transfected with WT or mutant mCherry-flSTIM1 and Orai1-GFP. 24–28 h after transfection, cells were loaded with 1 µM fura-2/AM (Invitrogen) in culture medium for 30 min at room temperature, washed, and plated on poly-D-lysine-treated coverslip chambers.

Ca^2+^ imaging was conducted as described previously (van Dorp et al., 2021) using a Zeiss 200M inverted microscope equipped with a Fluar 40X NA 1.3 oil-immersion objective. Cells were excited alternately at 350 and 380 nm (Polychrome II, TILL Photonics), and emission at 534 ± 30 nm (Semrock) was collected with a Flash4.0 sCMOS camera (Hamamatsu Corp.) with 2 × 2 binning. Images were background-corrected before calculating the mean 350/380 ratio for each cell. Standard 2 Ca Ringer’s solution contained (in mM): 155 NaCl, 4.5 KCl, 2 CaCl_2_, 1 MgCl_2_, 10 D-glucose and 5 Na-HEPES (pH 7.4). Ca^2+^-free (0 Ca) Ringer’s was prepared by replacing CaCl_2_ with 2 mM MgCl_2_ and 1 mM EGTA. Thapsigargin (Sigma-Aldrich) was diluted from a 1 mM stock in DMSO in Ca^2+^-free Ringer’s.

### Cysteine crosslinking and Western blot analysis

For crosslinking flSTIM1 in situ, STIM1/2 double knockout (DKO) HEK293 cells (Zhou et al., 2018) were co-transfected with mCh-STIM1 and HA-STIM1 constructs with selected residues replaced by cysteine. 48 h after transfection, cells were exposed to 0.2 mM diamide for 10 min in Ringer’s solution containing 2 mM Ca^2+^. Cells were then lysed in RIPA buffer containing 20 mM NEM, 0.5 mM EDTA, and protease inhibitor cocktail (Cell Signaling Technology). Samples were run on SDS-PAGE and analyzed by Western blot using mouse anti-HA antibody (1:2000, Sigma-Aldrich) and rabbit anti-mCherry antibody (1:2000, OriGene Technologies), with secondary antibodies (680RD anti-mouse and 800CW anti-rabbit) on a LI-COR Odyssey imaging system.

Western blots were scanned and analyzed using ImageJ, and the area under each peak indicated the amount of protein in each band. Crosslinking efficiencies were calculated by dividing the amount of STIM1 in the mCherry x HA heterodimer band (crosslinked) by an estimate of the total amount of STIM1 heterodimer present in each sample (crosslinked + non-crosslinked). The intensities of the HA and mCherry channels were normalized using the mCherry x HA heterodimer band, which contains the same amount of STIM1 in both channels. The total amount of heterodimer was estimated from the total amounts of STIM1 in the mCherry and HA channels, assuming random assembly of the two subunits of the dimer.

## ACKNOWLEDGEMENTS

The authors thank Dr. Mohamed Trebak (Univ. of Pittsburgh) for kindly supplying the STIM1/STIM2 double knockout HEK293 cell line, and members of the Lewis lab for helpful discussions. This work was supported by NIH grant R37GM45374, R35GM149305, and the Mathers Charitable Foundation (RSL).

## Notes

### Competing Interest Statement

The authors have declared no competing interest.

