## Supplementary Figures and Tables for "Multiple weak brakes act in concert to regulate STIM1 and control store-operated calcium entry"

**SUPPLEMENTARY FIGURE LEGENDS**

**Supplementary Figure 1. Comparison of SOCE evoked by WT STIM1 vs cysteineless STIM1**

STIM1/2 DKO HEK293 cells were cotransfected with Orai1 and either WT mCh-STIM1 or mCh-STIM1 with all endogenous cysteines (C49, C56, C227, and C437) replaced by serines ('cysless'). Fura-2 350/380 ratios are shown (mean ± sem; WT, from Fig. 5C, n=57 cells; cysless, n=54 cells). Solution changes are shown with [$Ca^{2+}$] in mM and 1 µM TG added as indicated to deplete $Ca^{2+}$ stores. Cysless STIM1 is fully functional as assessed by SOCE observed upon $Ca^{2+}$ readdition.

**Supplementary Figure 2. smFRET amplitude histograms for flSTIM1 in 2 mM $Ca^{2+}$**

Predominant peak FRET values and number of molecules used to construct the histogram are listed in Supplementary Table 1. Abbreviations: PC (phosphatidylcholine liposomes), 4 PC : 1 PS (4:1 PC and phosphatidylserine liposomes), 4EA (E318A/E319A/E320A/E322A), 4KA (K382A/K384A/K385A/K386A), 4S (L321S/V324S/L328S/L335S), TM-CT (STIM1 208-685), MBP-TM-CT (MBP+STIM1 201-685).

**Supplementary Figure 3. AlphaFold2 models of flSTIM1**

Top five models generated by AlphaFold2, using STIM1 residues 35-444 as input. For each model, the regions from the cEF hand through CAD (63-444) are shown. Domains are colored as shown in Fig. 1, and predicted membrane edges based on the TM domain are shown by dashed lines. Only model 2 is consistent with distances derived from smFRET.

**Supplementary Figure 4. Quantifying the resting state probability for flSTIM1, TM-CT, and MBP-TM-CT**

smFRET amplitude histograms from Fig. 2D are fitted with a sum of Gaussian curves to measure the fractional occupancy of the low-FRET peak for WT STIM1, TM-CT, and MBP-TM-CT.

**Supplementary Figure 5. Helical alignment of L261C in the CC1α1-CC3 brake**

A. Proximity of L261 (CC1α1) and T420 and L423 (CC3) predicted by the AlphaFold2 model.

B. Western blot showing heterodimer disulfide crosslinking between HA-STIM1-L261C and mCh-STIM1-T420C and -L423C. Each lane contains the lysate from diamide-treated cells coexpressing HA-STIM1-L261C and a single mCherry-STIM1 mutant (red) (see Methods).

C. Percent of heterodimers forming disulfide crosslinks, measured from the gel in B (see Methods). Results are representative of two experiments.

**Supplementary Figure 6. Effects of the 4EA mutation on stability of the resting STIM1 conformation**

A. smFRET histograms of 309:309' for WT (n=76) and 4EA (E318A/E319A/E320A/E322A) STIM1 (n=225) in 2 mM $Ca^{2+}$.

B. smFRET histograms of 242:242' for WT (n=361) and 4EA STIM1 (n=225) in 2 mM $Ca^{2+}$.

C. Fura-2 350/380 ratios (mean ± sem) from HEK cells expressing Orai1 and either STIM1-WT (from Fig. 5C, n=57cells), or STIM1-4EA (n=62 cells). Changes in extracellular [$Ca^{2+}$] (0 and 2 mM) and addition of thapsigargin (TG; 1 µM) are indicated. Resting $[Ca^{2+}]_i$ is elevated in cells expressing STIM1-4EA, indicating constitutive activity of STIM1 and SOCE, while TG-induced SOCE is normal.

**Supplementary Figure 7. Conformational dynamics of the flSTIM1 EF-SAM domain**

A. Probability distribution of the mean smFRET transition frequency over a 5-s period for 64:178 dye pair (EF-SAM; n=213).

B. smFRET histogram for 64:178 (top) and representative single molecule traces (below) showing stable and fluctuating molecules. Dashed lines indicate peak values from the smFRET histogram.

C. Probability distribution of the mean smFRET transition frequency over a 5-s period for the 178:178' dye pair (SAM-SAM; n=271).

D. smFRET histogram for 178:178' (top) and representative single molecule traces showing stable and fluctuating molecules. Dashed lines indicate peak values from the smFRET histogram.

### SUPPLEMENTARY TABLES

**Supplementary Table 1. Summary of smFRET measurements for the resting state of flSTIM1 in saturating (2 mM) $Ca^{2+}$**

For each dye pair, the predominant smFRET value from the amplitude histograms in Supplementary Fig. 2 is listed and used to estimate a distance. The number of single molecule traces used to construct each histogram is indicated.

**Supplementary Table 2. Comparison of smFRET measurements in flSTIM1 vs ctSTIM1**

The table lists the predominant smFRET values for each indicated dye pair. ctSTIM1 values are reproduced from our previous study (van Dorp et al., 2021).

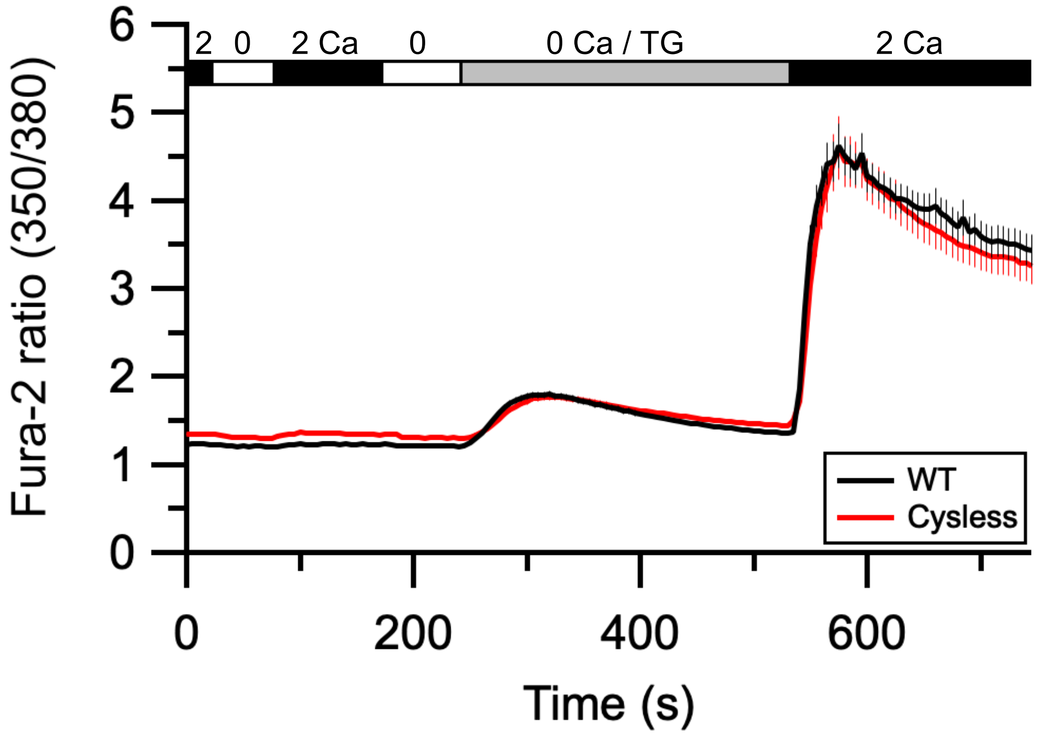


**Suppl. Figure 1**

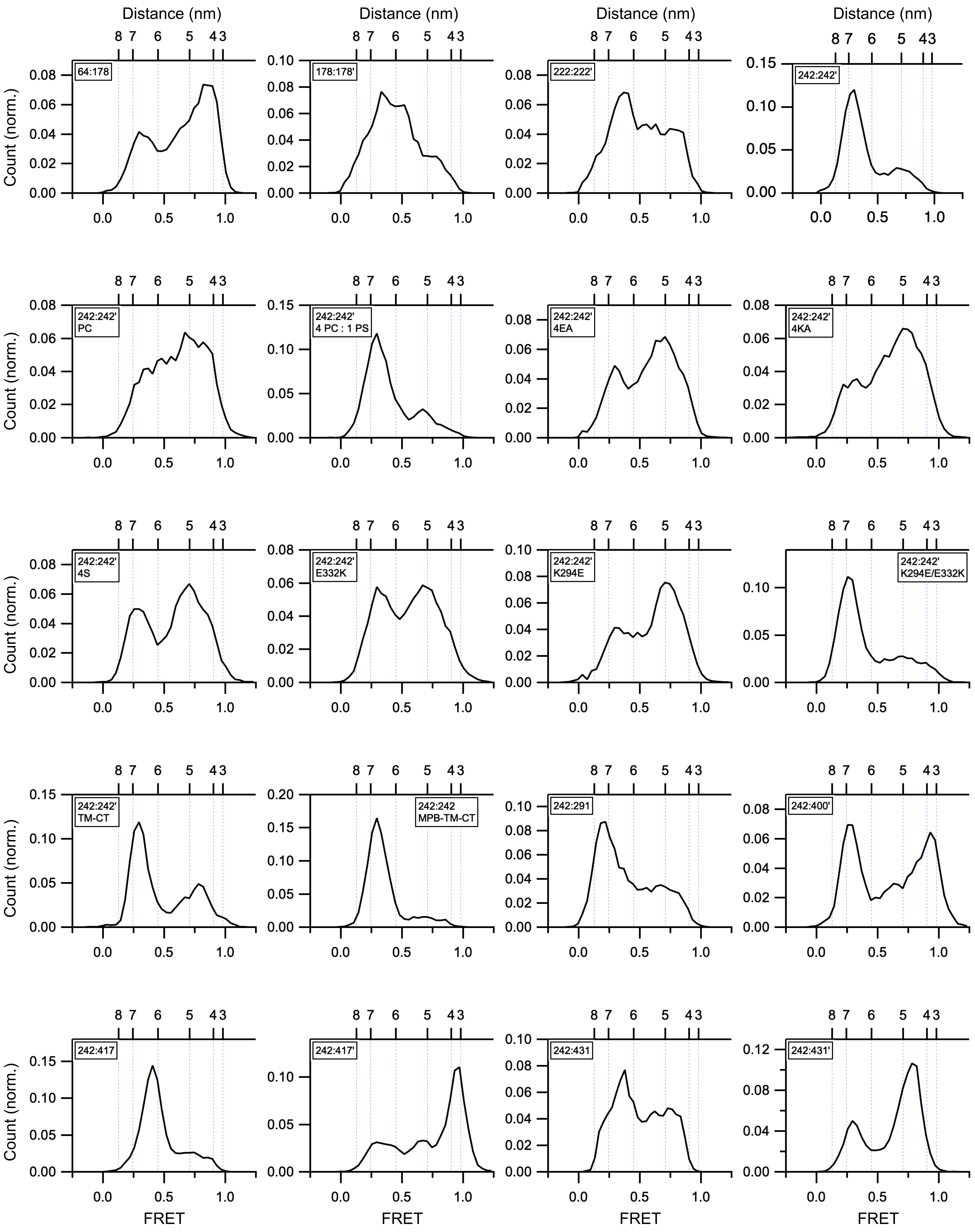


**Suppl. Figure 2 (pt 1/2)**

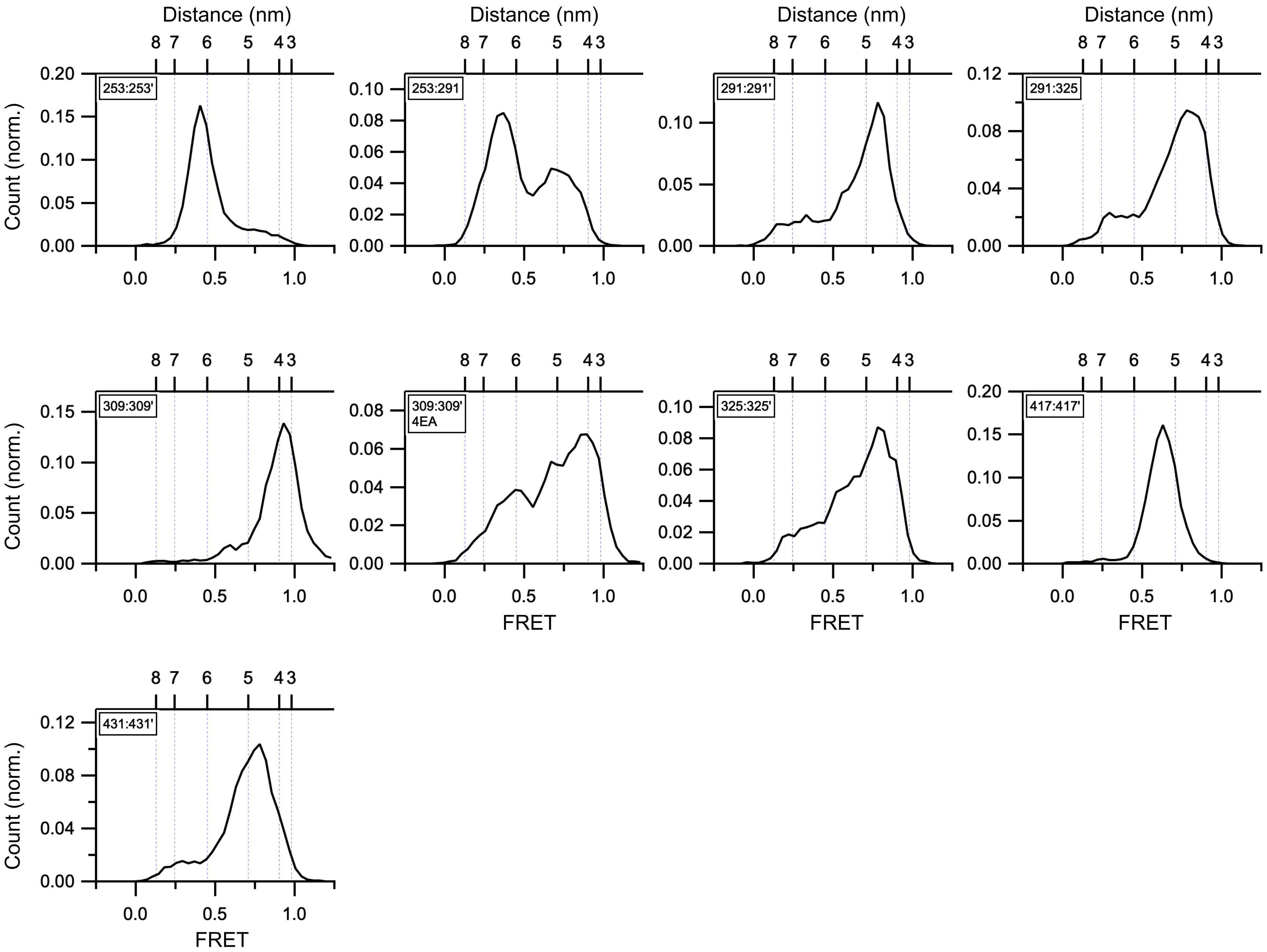


**Suppl. Figure 2 (pt 2/2)**

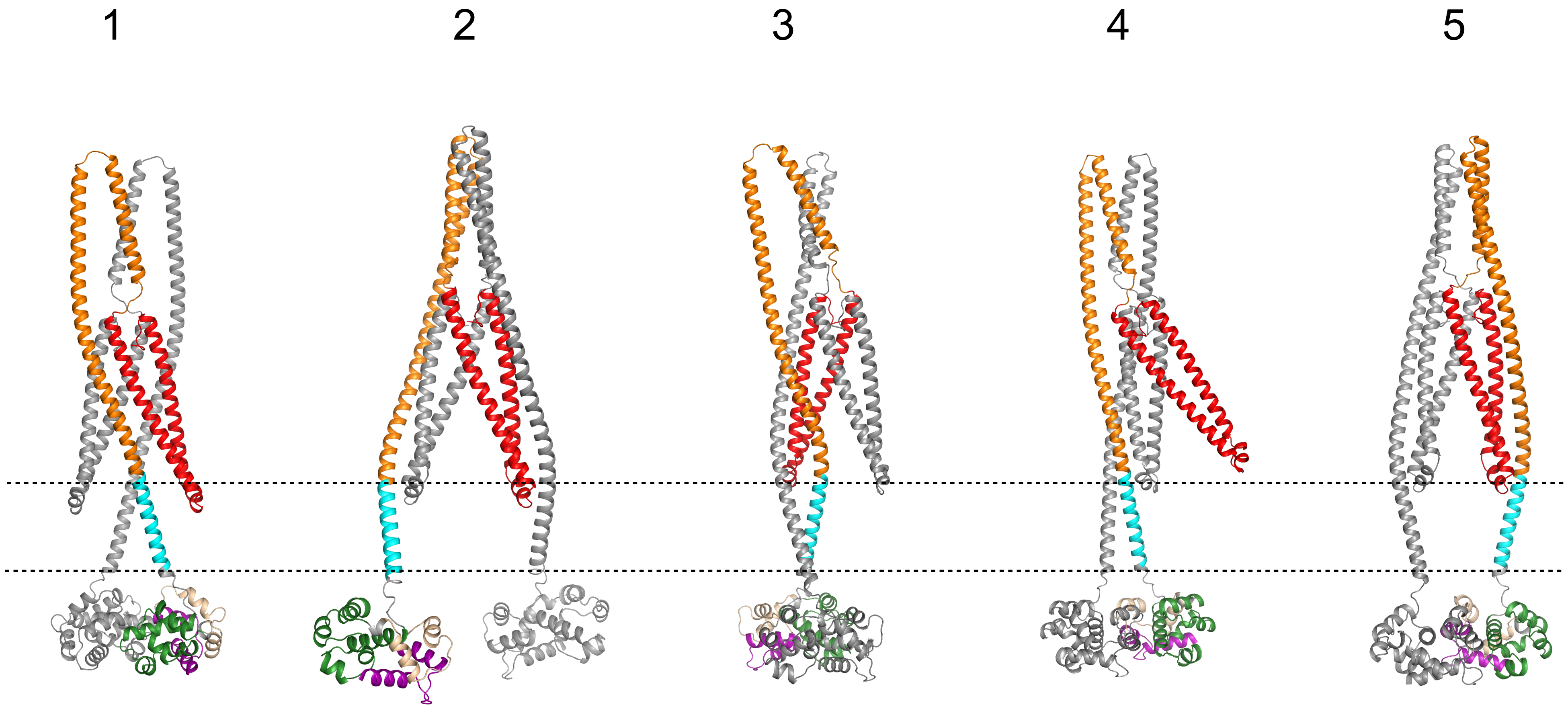


**Suppl. Figure 3**

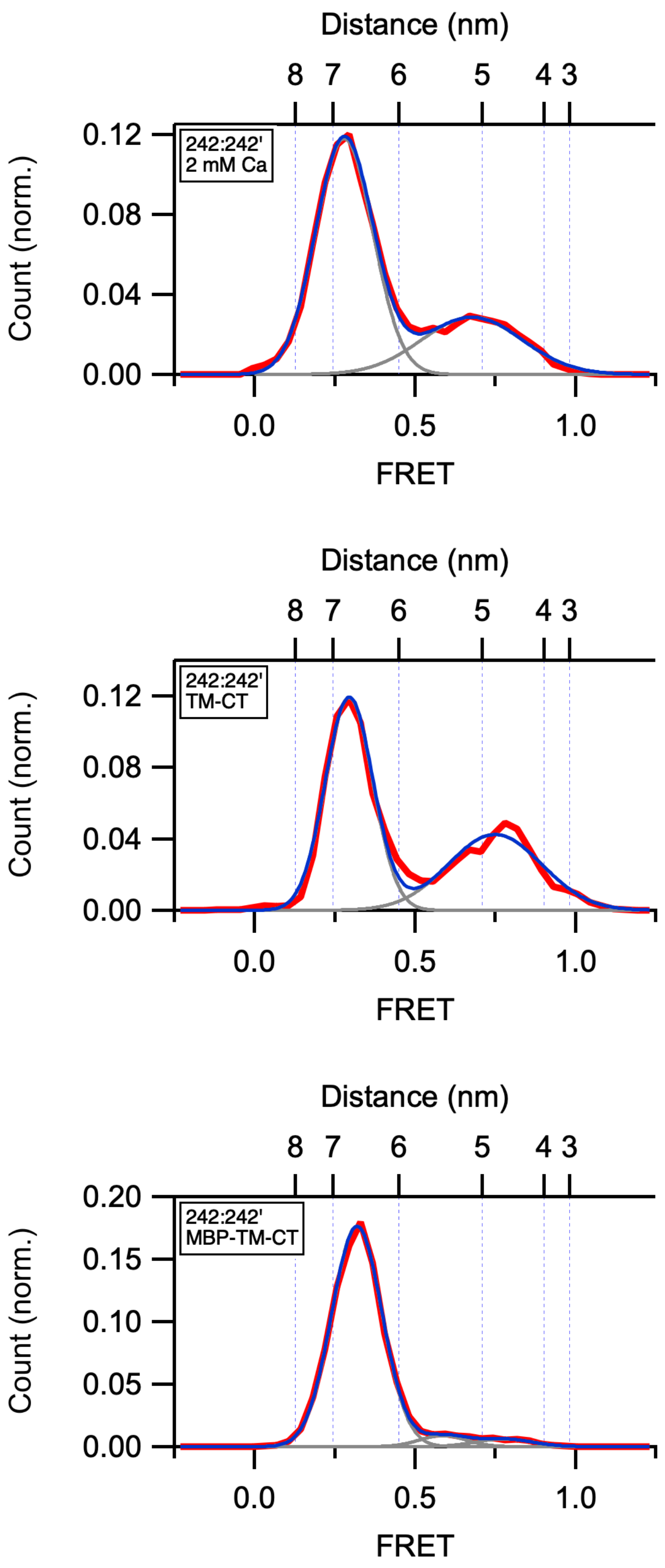


**Suppl. Figure 4**

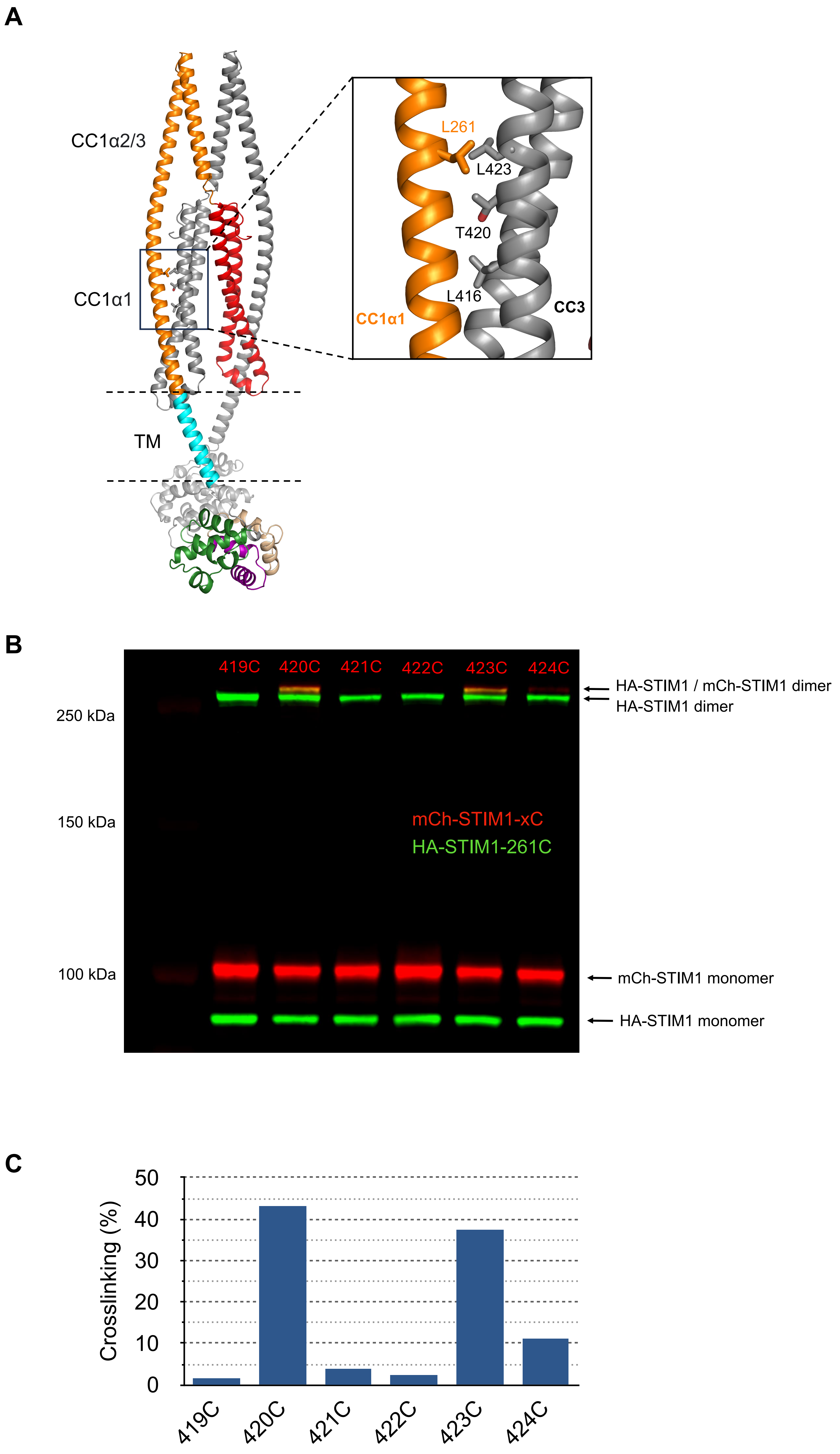


**Suppl. Figure 5**

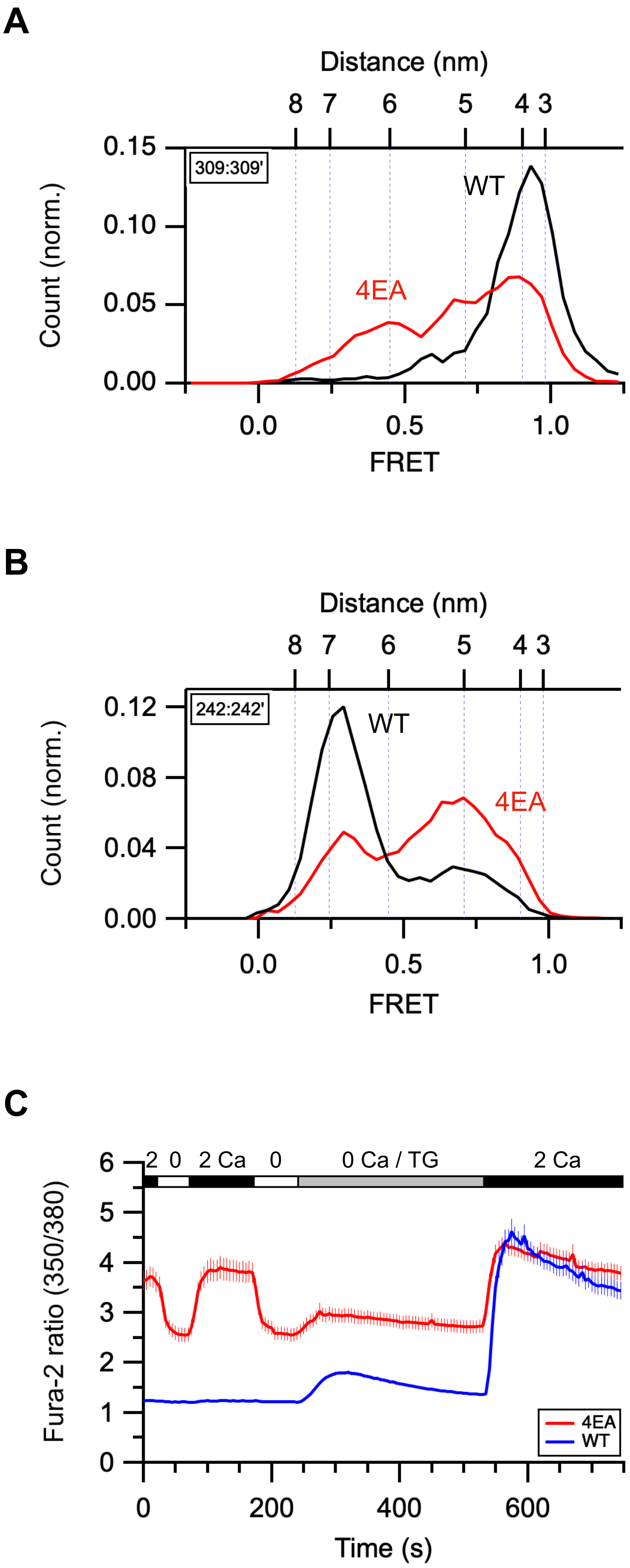


**Suppl. Figure 6**

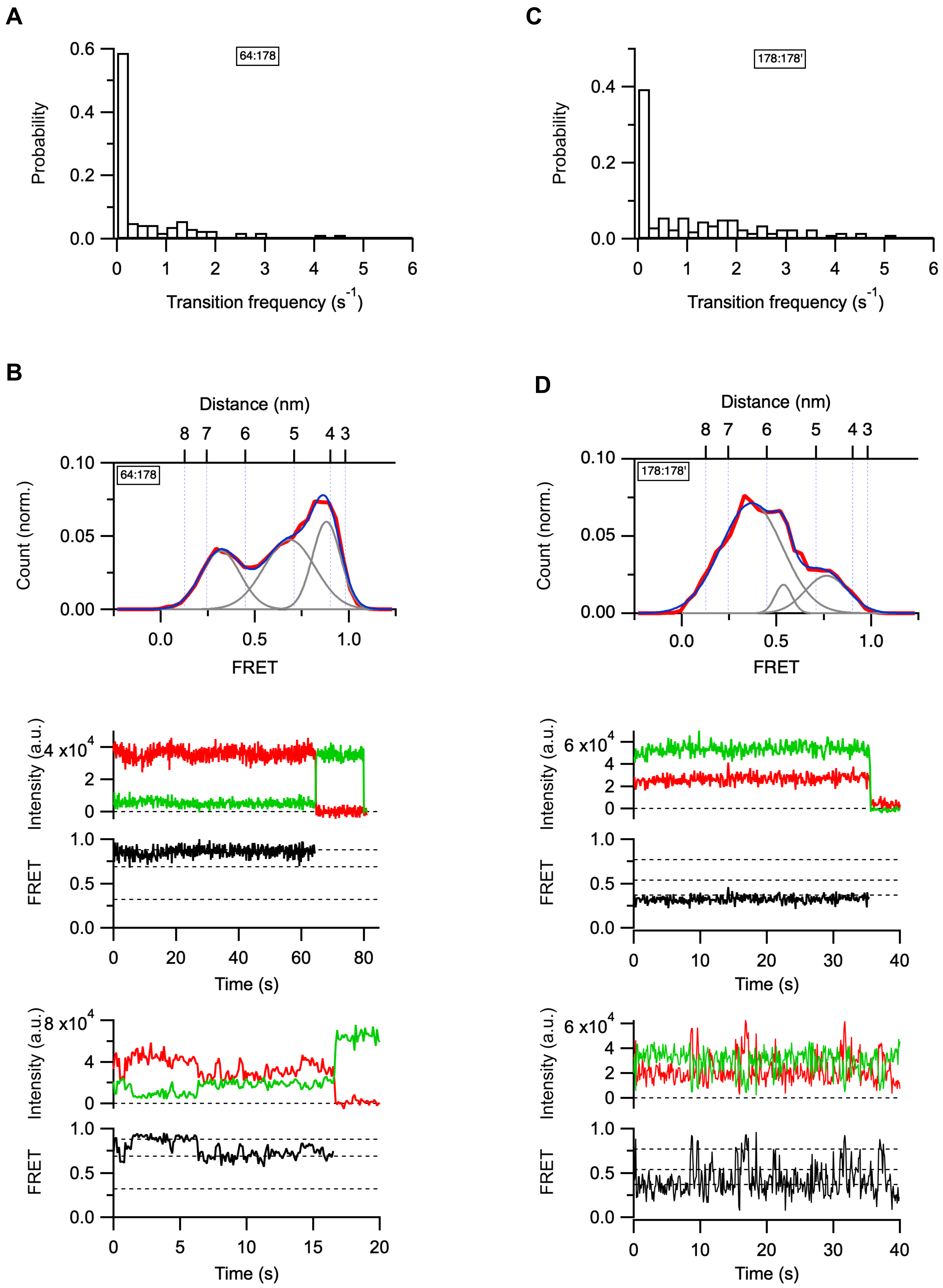


**Suppl. Figure 7**

| Dye pair | FRET | Distance (nm) | Number of molecules (n) |
|---|---|---|---|
| 64:178 | 0.84 | 4.40 | 213 |
| 178:178' | 0.35 | 6.43 | 271 |
| 222:222' | 0.35 | 6.43 | 114 |
| 242:242' | 0.28 | 6.79 | 361 |
| 242:291 | 0.22 | 7.16 | 192 |
| 242:400' | 0.94 | 3.67 | 188 |
| 242:417 | 0.4 | 6.21 | 291 |
| 242:417' | 0.96 | 3.42 | 190 |
| 242:431 | 0.34 | 6.48 | 298 |
| 242:431' | 0.79 | 4.65 | 189 |
| 253:253' | 0.43 | 6.08 | 281 |
| 253:291 | 0.38 | 6.29 | 254 |
| 291:291' | 0.8 | 4.60 | 185 |
| 291:325 | 0.8 | 4.60 | 202 |
| 309:309' | 0.94 | 3.67 | 76 |
| 325:325' | 0.81 | 4.55 | 243 |
| 417:417' | 0.63 | 5.31 | 169 |
| 431:431' | 0.8 | 4.60 | 204 |

**Suppl. Table 1**

| Dye pair | flSTIM1 | ctSTIM1 |
|---|---|---|
| 242:242’ | 0.27 | 0.31 |
| 242:400’ | 0.94 | 0.95 |
| 242:417 | 0.4 | 0.4 |
| 242:417’ | 0.96 | 0.9 |
| 242:431 | 0.34 | 0.3 |
| 242:431’ | 0.79 | 0.73 |
| 309:309’ | 0.93 | 0.9 |
| 417:417’ | 0.63 | 0.64 |
| 431:431’ | 0.79 | 0.77 |

**Suppl. Table 2**
